# uPIC–M: efficient and scalable preparation of clonal single mutant libraries for high-throughput protein biochemistry

**DOI:** 10.1101/2021.08.04.455146

**Authors:** Mason J. Appel, Scott A. Longwell, Maurizio Morri, Norma Neff, Daniel Herschlag, Polly M. Fordyce

## Abstract

New high-throughput biochemistry techniques complement selection-based approaches and provide quantitative kinetic and thermodynamic data for thousands of protein variants in parallel. With these advances, library generation rather than data collection has become rate limiting. Unlike pooled selection approaches, high-throughput biochemistry requires mutant libraries in which individual sequences are rationally designed, efficiently recovered, sequence-validated, and separated from one another, but current strategies are unable to produce these libraries at the needed scale and specificity at reasonable cost. Here, we present a scalable, rapid, and inexpensive approach for creating <u>U</u>ser-designed <u>P</u>hysically <u>I</u>solated <u>C</u>lonal–<u>M</u>utant (uPIC–M) libraries that utilizes recent advances in oligo synthesis, high-throughput sample preparation, and next-generation sequencing. To demonstrate uPIC–M, we created a scanning mutant library of SpAP, a 541 amino acid alkaline phosphatase, and recovered 94% of desired mutants in a single iteration. uPIC–M uses commonly available equipment and freely downloadable custom software and can produce a 5000 mutant library at 1/3 the cost and 1/5 the time of traditional techniques.

## INTRODUCTION

Recent technological advances enable the biochemical interrogation of many protein variants in parallel with the precision and versatility needed to dissect mechanisms of function. These techniques, termed broadly here as *high-throughput biochemistry* (HTB), report *quantitative* kinetic and thermodynamic measurements for thousands of individual protein sequences. This advance is made possible by developments in programmable automated liquid handling that increase the scale of plate-based assays,^1,2^ and recently, by a miniaturized microfluidic platform that allows parallel measurement of thousands of variants on one microscope slide.^3–5^ For basic enzymology and biophysics studies, HTB approaches using mutational scanning libraries allow identification of the effects of all residues on folding, stability, binding and catalysis. To advance precision medicine, libraries comprised of human allelic variants^6^ can be assayed for folding and function and “variants of uncertain significance” can be classified by their biophysical propensity to drive disease or respond to therapeutics.^7,8^ Finally, within evolutionary biology, measurements of many extant orthologs and ancestral reconstructions can elucidate the molecular underpinnings of evolutionary adaptation (Figure 1A).^9–12^ Each of these applications requires moving beyond simply identifying mutants with desired properties from large-scale screens to directly linking each of many sequence perturbations with its functional effects.

**Figure 1.**
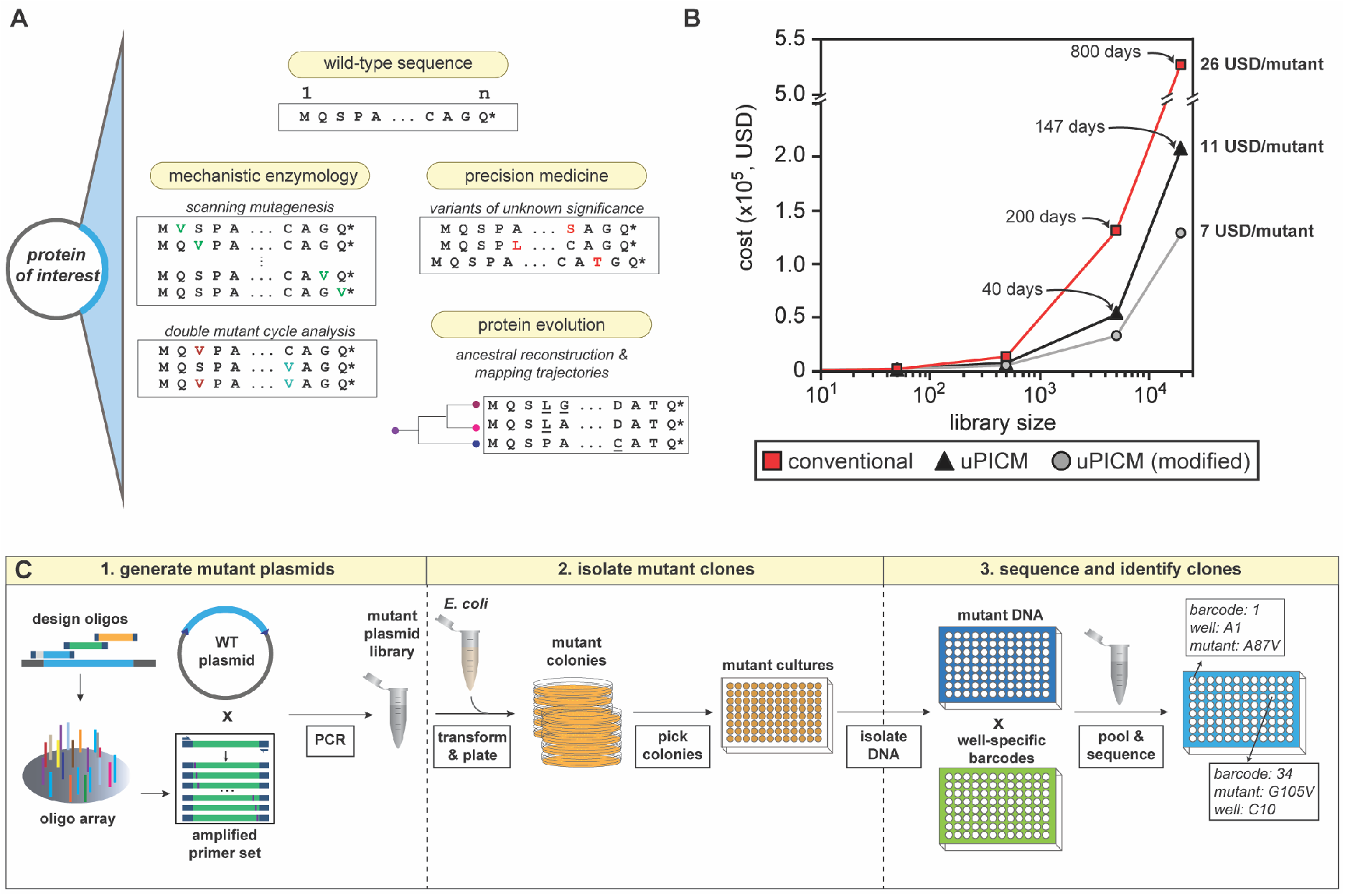
Overview of uPIC–M pipeline to generate user-defined clonal mutant libraries. (**A**) Examples of clonal libraries from uPIC–M and potential high-throughput biochemistry applications. Applications are listed along with examples of the types of variants involved. (**B**) Comparison of cost (including materials and labor) of conventional mutagenesis vs. uPIC–M for libraries of 50–20000 mutants. A uPIC–M clone sampling rate of 384 per 50 desired mutants (7.68-fold excess) was used for these calculations. uPIC–M (modified) represents a lower cost version of uPIC–M with the addition of pipet tip washing for plate liquid transfer steps. (**C**) Workflow for generating uPIC–M libraries in three phases: (1) Mutagenic oligos are synthesized for ∼50 residue windows on a pooled array and selective PCR amplification of each window generates a primer pool used for QuikChange; (2) pooled QuikChange reactions are transformed and plated, with each plate containing a mixture of ∼50 possible single mutants, facilitating colony picking into multiwell plates to isolate clonal libraries of unidentified variants; (3) clonal libraries are prepared and sequenced by NGS to reveal the genotype and location of each variant.

As high-throughput biochemistry tools increase the throughput of quantitative protein measurements by 10^2^–10^3^-fold,^1–5^ generating the requisite variant libraries has emerged as the new bottleneck. For HTB to provide measurements for rationally chosen protein variants, input libraries must be user-defined clonal mutant libraries in which individual mutants are sequence-validated and physically isolated from one another for downstream assays.

Conventional site-directed mutagenesis generates user-defined, isolated variants by performing each mutagenesis reaction, plasmid isolation step, and downstream sequencing within physically separated reactions. This approach results in high *control* (the ability to create only mutants of interest), but is prohibitively costly and labor-intensive for applications requiring >100 variants (Figure 1B). Conversely, existing techniques for generating mutant libraries, while powerful, are typically not suited for generating large-scale user-defined clonal mutant libraries.^13^ For example, error-prone PCR^14–16^ and mutagenic oligos containing degenerate codons^17,18^ allow generation of extremely large mutant libraries (10^7^–10^9^) at relatively low cost; such libraries are ideal for selecting constructs with desired characteristics, but these mutagenesis strategies do not allow generation of a desired set of defined sequences.

Here, we introduce uPIC–M (<u>U</u>ser*-*designed <u>P</u>hysically <u>I</u>solated <u>C</u>lonal–<u>M</u>utant) libraries, a method to prepare the needed mutant libraries that dramatically reduces the time and cost of conventional mutagenesis to empower high-throughput biochemistry (Figure 1C). uPIC–M can create 10^2^–10^4^ mutants at a material and labor cost of ∼$11 USD/mutant in 40 days for 5000 mutants, compared to an estimated $26 USD/mutant in 200 days for conventional mutagenesis (Figure 1B). The uPIC–M pipeline includes three stages of library production: (i) user-directed pooled mutagenesis, using commercially-available oligo arrays and a simplified in-house oligo design tool; (ii) isolation of mutant clones with widely-available robotic pickers; and (iii) next-generation sequencing (NGS) to identify clone sequences and their locations, leveraging recent automation developments from single-cell sequencing.^19^ uPIC–M uses the robust and accessible Illumina sequencing platform and a combination of existing open-source and custom analyses available on public software repositories to rapidly identify and evaluate library variants.

To develop and test uPIC–M, we set out to produce a scanning mutant library encoding single substitutions for every position in a 541 amino acid enzyme. Guided by stochastic sampling simulations, we picked a total of 4992 colonies to yield 3530 fully sequenced clones containing 507 desired single alanine and valine mutants, representing a library coverage of 94%. The efficiency and speed of this platform will accelerate the adoption and expand the scope of HTB.

## RESULTS AND DISCUSSION

### Overview of uPIC–M

The uPIC–M library generation pipeline consists of three stages (Figure 1C, 1–3) over approximately 8 days (Figure S1). During stage 1 (“generate mutant plasmids”), *E. coli* are transformed with pooled libraries of mutant plasmids generated via QuikChange-HT mutagenesis using user-defined, array-synthesized mutagenic oligonucleotides to create the specified variants. During stage 2 (“isolate mutant clones”), transformed *E. coli* are plated to isolate individual mutant colonies, which are then picked and used to inoculate liquid cultures within multiwell plates. During stage 3 (“sequence and identify clones”), mutant DNA is amplified and “barcoded” with well-specific primer sequences (“barcodes”) prior to pooling for NGS. For amplicons longer than 600 nucleotides (the maximum read length of typical paired end Illumina sequencing reads), amplified sequences can be fragmented using Tn5 transposase prior to barcoding to ensure the ability to acquire and associate reads spanning the complete amplicon. This barcoding strategy allows parallel sequencing while i) preserving the plate-well origin of each read, and ii) providing a means to group reads for reconstructing the full-length sequence of each clone.

After sequencing, NGS reads are first demultiplexed according to the library barcode (here: 4992 barcodes); reads are then grouped by the barcodes specifying each well and aligned to the WT “reference” amplicon sequence and variants are “called” from these aligned sequences. uPIC–M thus reports the full-length ORF sequence, physical well location, and quality information of clonal library variants, allowing users to select thousands or more single mutant clones of interest to create curated libraries for downstream high-throughput biochemistry applications.

### Design of tiling ORF windows allows selective mutagenesis from oligo arrays

We used QuikChange-HT mutagenesis, an oligo array-based strategy that provides rationally chosen mutants and offers the following advantages: 1) a simple experimental procedure, thus increasing throughput; 2) the ability to selectively amplify distinct mutagenic oligo subsets from the same array, permitting the use of the same source array for different experiments and targets; and 3) the ability to implement a design strategy that disfavors the production of double and higher-order mutants during pooled mutagenesis reactions, reducing otherwise-costly downstream sampling of clones to identify the desired single mutants.^20^ Other previously reported methods can generate large libraries of rationally chosen mutants from oligo arrays but lack these time- and cost-saving features.^21,22^

QuikChange-HT generates mutants by a straightforward approach, the same as conventional PCR mutagenesis, but uses a unique mutagenic oligo design strategy that meets our needs. Coding regions are first divided into ∼200–300 nucleotide “windows” (with the exact length dependent on maximum oligonucleotide synthesis length and cost/nt). The 5’ and 3’ termini of each window (∼25 nt each) act as universal primer sites for amplification of that window from the pooled arrays and the ∼150 nt intervening sequence carries user-defined codon substitutions across ∼50 residues that will be introduced by QuikChange (Figure 2A). Overlapping adjacent windows by ∼20–30 bp makes it possible to uniquely amplify all mutagenic oligonucleotides within single windows with a corresponding primer pair (Figure S2). This strategy allows mutagenic oligos for many uPIC– M targets to be encoded by the same parent array, greatly reducing the cost (per oligo) and makes it possible to continue to generate mutagenic oligos from the array via PCR. Downstream mutagenesis reactions use the amplified mutagenic oligos as primers to produce pooled mutant sublibraries, and proceed by iteratively denaturing double-stranded plasmid DNA, annealing the oligonucleotide that encodes the desired mutation, and extending via a high-fidelity polymerase. After rounds of annealing and extension, parental methylated and hemi-methylated strands are digested via DpnI prior to transformation. The window approach reduces the likelihood of double and higher-order mutants by dividing sublibraries into separate reactions, which contain only mixtures of mutagenic primers that share the same termini sequences. As such, pooled mutagenic primers bind competitively to the same sequence of template DNA..

**Figure 2.**
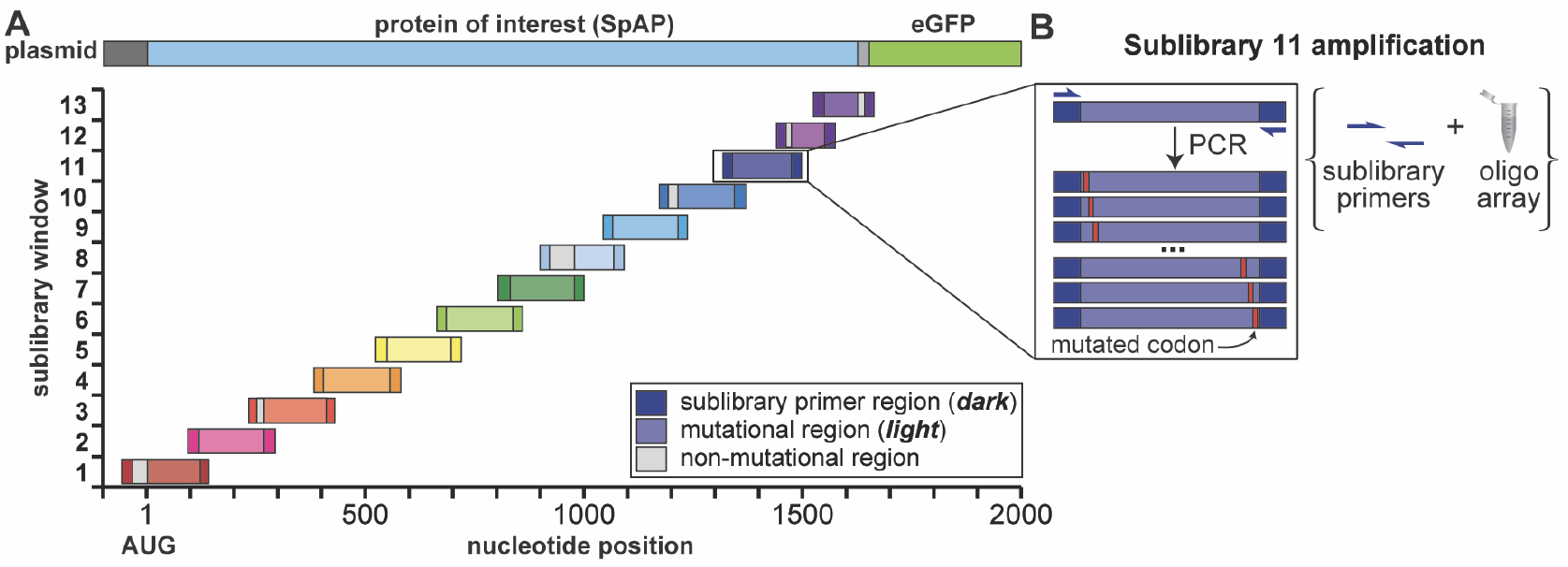
Tiling window strategy for uPIC–M mutagenic oligo array design. (**A**) A tiling window strategy (see Figure 1C) divides the ORF from the protein of interest into mutagenic sublibrary regions, with sublibrary oligo length constrained by DNA synthesis limits. Each window contains unique forward and reverse priming sites (dark shading, here ∼25 nt each) at the 5’- and 3’-termini surrounding a mutational region (light shading, here ∼150 nt). For a scanning library, each codon along the length of a sublibrary mutational region is substituted via an individual mutagenic oligo. (**B**) Selective amplification of oligos from a single window (Sublibrary 11). Forward and reverse primers specific to a single sublibrary are used to amplify oligos from the resuspended array material, yielding an oligo pool containing ∼50 codon substitutions from the same mutagenic window.

To develop and demonstrate the capabilities of uPIC–M, we designed a library of mutagenic primers to mutate each residue of the 541 amino acid alkaline phosphatase SpAP to Ala or Val. This design process generated 13 mutational windows to efficiently encode the selected valine or alanine substitution at each position (Table S1).

### QuikChange-HT mutagenesis

Subsets of mutants are created in sublibrary pools, with one mutagenesis reaction carried out per sublibrary. To generate the material for each of 13 mutagenesis reactions for SpAP, we first amplified mutagenic primers for a given “window” from the total oligonucleotide pool via PCR and window-specific primers (Figure 2B, Table S1). Following spin-column purification, these amplified primers were used directly as QuikChange-HT mutagenic primers. Agilent-designed (see Materials and Methods) primers resulted in clean amplification of sublibrary mutagenic primer pools (Figure S2) from an array containing scans for SpAP as well as four additional genes (see data repository for full array sequence) with purified yields of ∼14–50 nM each (Table S2). We then performed mutagenesis reactions for each sublibrary following standard QuikChange protocols (linear PCR amplification of WT template followed by DpnI digestion).

### Simulated mutant sampling to predict screening requirements

For randomly sampled clones from a pool of variants, one needs more than the number of desired mutants to obtain complete or near-complete sampling, as the probability of obtaining a *novel* variant (one that has not already been sampled) decreases with increased sampling (similar to “the birthday problem” or the related “coupon collector’s problem” in probability theory). To estimate the number of clones that must be sampled to recover a given fraction of mutants from a specified variant population, we simulated stochastic sampling experiments in which we sampled clones *N* times from a pool of *M* variants without replacement. For libraries of 50, 500, and 5000 variants: 110, 1150, and 11600 draws were required to recover ≥90% of desired clones, respectively (Table S3). To consider how the presence of WT clones or unwanted variants (*e*.*g*. undesired single mutants and/or higher-order mutants) affect recovery rates, an additional term was added specifying the probability that any given draw returns a single mutant (Figure 3A). As expected, lower rates of single mutant recovery led to a requirement for more clone sampling to obtain equivalent library coverage (Table S3).

**Figure 3.**
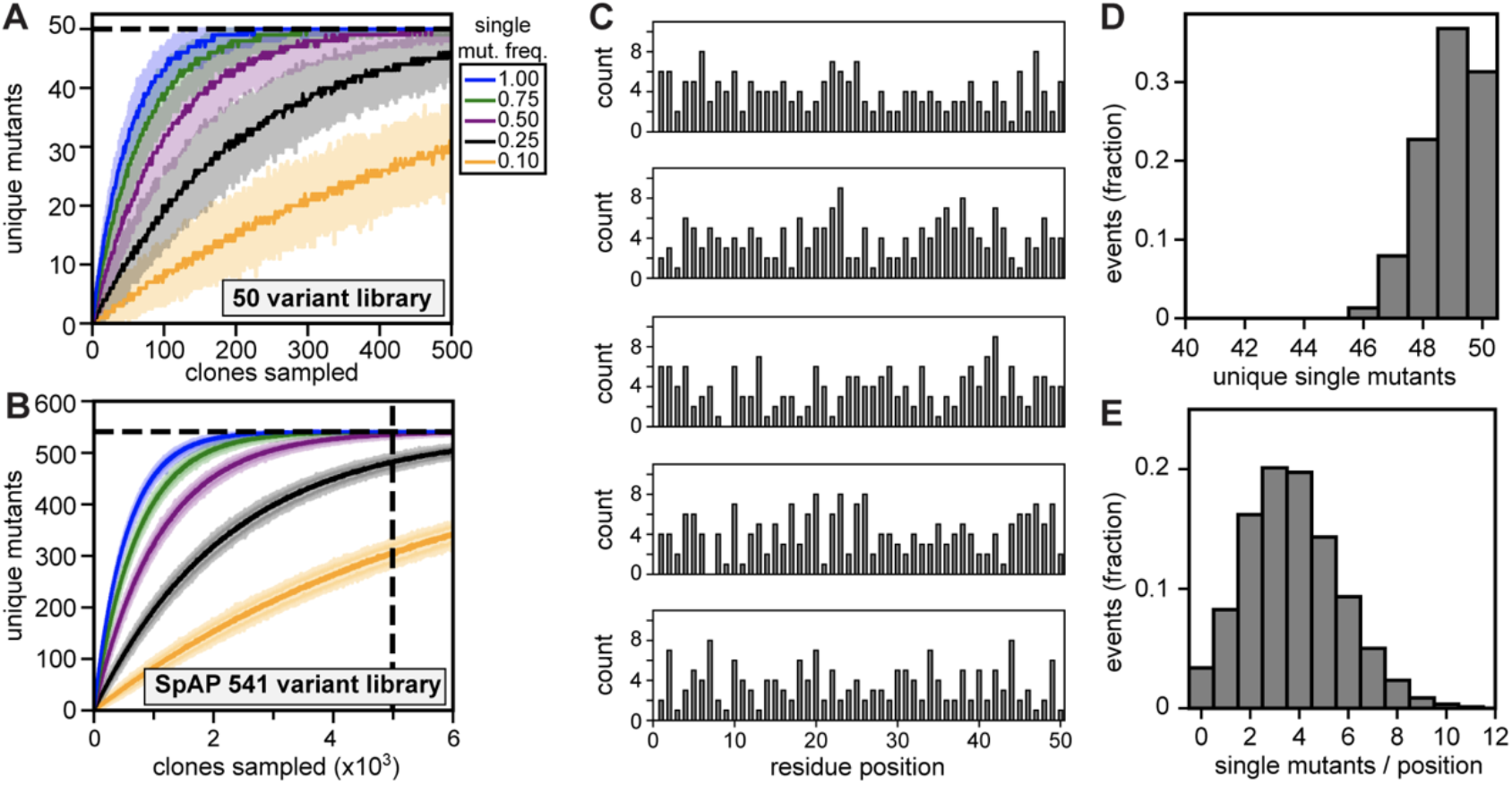
Simulated sampling of pooled single mutant libraries. (**A, B**) Simulation of the number of unique mutants obtained as a function of the number of clones sampled for pooled libraries containing 50 (**A**) or 541 (**B**) unique single mutants with single mutant frequencies from 0.1–1.0. The remaining fraction of each pool represents all other variants (*e*.*g*., WT, indels, double, and higher-order mutants). Each curve represents the average of 10^3^ simulations; shaded bands represent the 95% confidence interval; horizontal dashed lines (**A, B**) indicate the total possible number of unique mutants; vertical line (**B**) indicates the number of colonies picked for the SpAP library constructed herein (for legend, see **A**). (**C-E**) Simulated picking results for a sublibrary containing 50 single mutants at equal relative abundances sampled 384 times with a single mutant frequency of 0.5. (**C**) Simulated positional frequencies of single mutants; the results of five sampling simulations were chosen at random. (**D**) Histogram of expected mutant yields and (**E**) histogram of expected yields per sublibrary position (from 10^3^ sampling events).

We used these stochastic simulations to estimate the number of clones required to recover ≥90% of desired mutants within the 541 amino acid scanning mutagenesis library (V and A substitutions) for the SpAP construct (Figure 3B). To estimate the rates at which QuikChange-HT mutagenesis returns desired single mutants, we performed a preliminary pooled mutagenesis reaction, plated transformed *E. coli*, and Sanger sequenced 96 isolated clones. This preliminary sampling experiment returned 11 WT, 60 single mutant, and 5 double, triple and greater mutant constructs, and 20 additional clones with indels and/or sequencing errors, suggesting an approximate single mutant rate of 63% (Table S4). We elected to oversample each sublibrary, with up to 384 possible clones for each set of up to 50 desired mutants. Simulating this sampling ratio with a single mutant rate of 50% for 50 possible mutants, predicts a 92–100% yield (46–50 mutants) (Figure 3C, D). The distribution of the expected number of mutants per position obtained from random sampling revealed expected distributions of 0–11 mutants recovered at each position with a median of 4 (95% confidence interval of 0–8) (Figure 3E). The SpAP sublibraries encoded variable numbers of single mutants (range 25–48 possible mutants each, Table S1). Sampling at an approximately 384:50 clone to mutant ratio is a compromise as the increase in time is negligible (e.g. compared to sampling half as many clones), and still results in substantial savings in costs compared to conventional mutagenesis (Figure 1B).

### Clonal mutant isolation from plasmid libraries by pick-and-grow step

Clones must be physically isolated, both as a requirement of downstream high-throughput biochemistry assays, and to permit sequence identification and validation (Figure 1C, (2)). To facilitate separation via robotic colony picking, we transformed chemically competent *E. coli* with pooled mutagenesis reactions for each sublibrary mutational window and then plated transformations on LB agar plates (150 mm) supplemented with antibiotic for outgrowth to saturation overnight at 37 °C. These reactions produced a range of colonies (25–440 colonies/plate, Table S5) despite the use of identical concentrations of WT template and sublibrary primer concentrations in each (15 nM stock concentrations). As robotic colony selection by imaging requires colonies within a narrow range of size, shape, and density (∼300–500 evenly spaced colonies per 150 mm plate), we replated sublibrary transformations that were outside of this range at higher or lower density (6 of 13); for reactions that still yielded insufficient densities (3 of 13), we successfully repeated QuikChange reactions at the highest stock sublibrary primer concentrations (Table S5). Guided by our stochastic sampling simulations, we selected ∼384 colonies for each sublibrary, with a throughput of 8–10 384 well plates/day, for a total of ∼1.5 days for the 13 SpAP sublibrary plates. The robotic colony picker occasionally picked at the interface of multiple colonies, likely leading to mixtures of multiple variants within some wells. For significantly larger mutant libraries, alternative robotic systems that allow automated agar source and multiwell destination plate handling or single microbe^23^ or single droplet-based^24^ methods for cell sorting could significantly enhance throughput.

### Preparation of mutant DNA amplicons

DNA derived from mutant plasmids in *E. coli* clones must be amplified and enriched prior to NGS library preparation (Figure 1C, (3)). We generated amplicons by PCR (instead of isolating plasmids), as PCR requires minimal sample handling and produces linearized products that are directly compatible with downstream steps (sequencing library preparation and cell-free expression for HTB assays; Figure 4A, C). We amplified a 2525 bp region from each clone using universal primers complementary to the 5’- and 3’-UTR regions surrounding the SpAP-eGFP coding sequence (see Materials and Methods). To reduce contamination from *E. coli* genomic DNA in the final library, we systematically diluted liquid culture templates and measured contamination by qPCR (Figure S3). A 1:1000 dilution of six sample mutant cultures into H_2_O (corresponding to a final dilution of 1:5000) reduced *E. coli* DNA contamination to the limit of detection. To generate uniform amounts of PCR product from variable amounts of DNA templates within culture plates, we performed 25 cycles of PCR using a high-fidelity polymerase (Figure S4). We selected these conditions for amplification of mutant DNA from SpAP sublibrary plates. After dilution and amplification, DNA concentrations measured for half of the wells within each 384 well plate varied by ∼5-fold across all samples, and with median concentrations of ∼20–60 ng/μL (Table S6, Figure S5).

**Figure 4.**
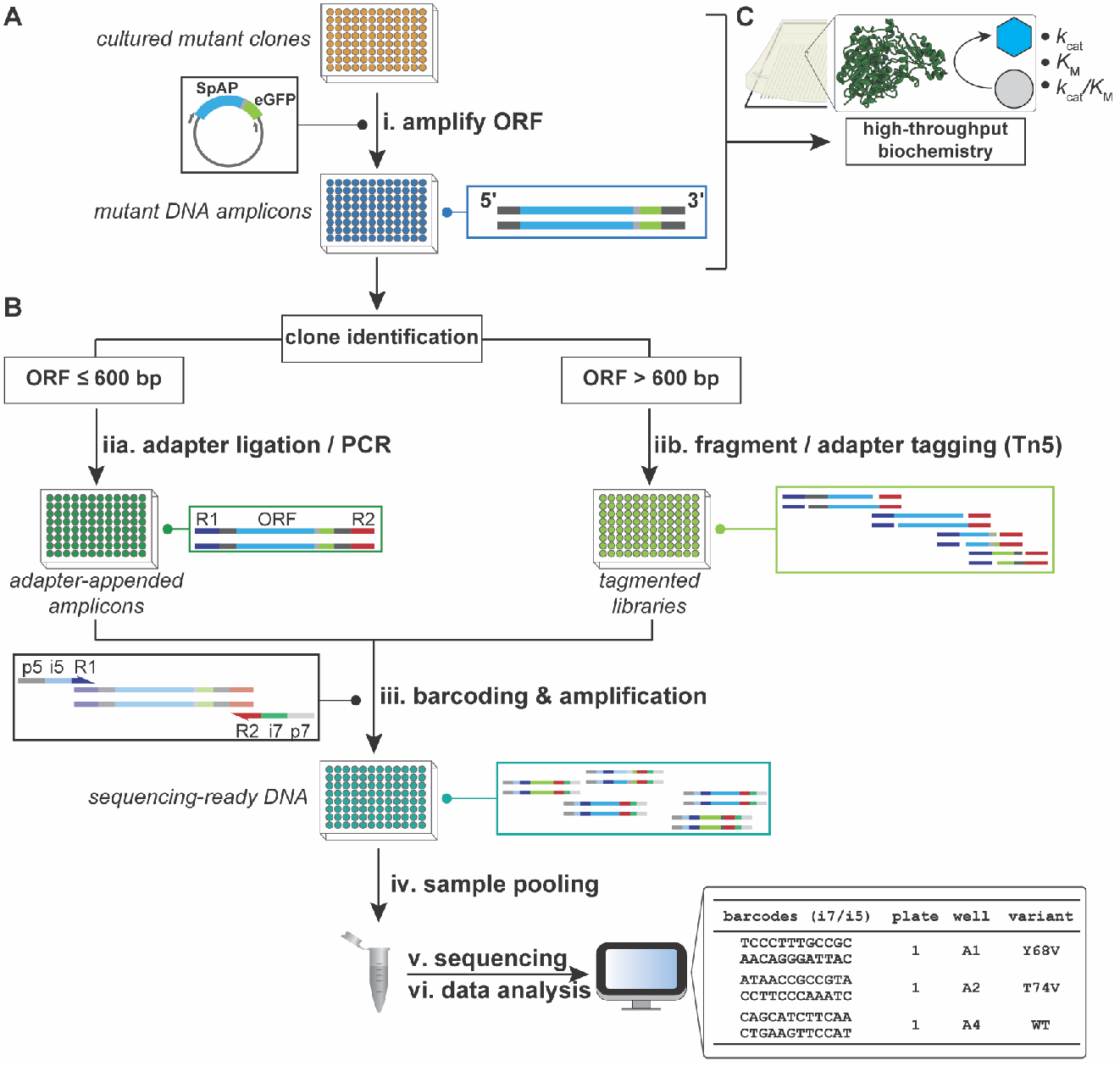
Schematic of uPIC–M sequencing library preparation. Preparation of sequencing libraries takes place in multi-well plate format (96 or 384) via the following steps: (**i**) ORF regions of target plasmids are amplified from each clone using universal primers to obtain enriched amplicon DNA; (**iia**) For amplicons ≤ 600 bp, universal Illumina adapters may be ligated directed to amplicons or added by amplification in a second PCR step; (**iib**) For amplicons >600 bp, DNA is fragmented and tagged using adapter-loaded Tn5 transposase, i.e. *tagmented*; (**iii**) amplicons or fragments are further amplified with Nextera primers that incorporate dual-unique i7 and i5 index barcodes; (**iv**-**vi**) amplified and barcoded clonal libraries are pooled for NGS, sequenced, and barcodes are used to report the plate-well location and genotype of each variant. (**C**) Mutant amplicons generated at (**i**) can be used directly for high-throughput biochemistry applications (shown here: cell-free expression and fluorogenic assay of an enzyme library using a microfluidic platform to obtain kinetic parameters).

### Tagmentation and barcoding of mutant amplicons

Mutational regions spanning <600 nucleotides can simply be barcoded and the entire region sequenced using 2 × 300 paired end Illumina reads (Figure 4B, iia).^25^ Longer ORFs, as are common and is the case for our example, require an alternative step to enzymatically fragment DNA and associate well-specific barcodes with each fragment, as used here (Figure 4B, iib). Critically, both strategies install universal adapter sequences to the DNA within each sample well, providing priming sites for barcodes that are specific to each well in a subsequent amplification step. Following Tn5 tagmentation of the 2525 bp SpAP-eGFP amplicons, we used the universal adapter sequences attached to fragment ends as priming sites to amplify DNA and add sequences required for Illumina sequencing, including: (1) sequences required to bind amplicons to sequencing flow cells (p5/p7), (2) plate/well-specific index 1 and index 2 barcodes (i7/i5), and (3) complementary sites for sequencing primers (R1 and R2) (Figure 4B, iii).^26,27^ All barcoded samples can then be pooled and sequenced in a single run via NGS (Figure 4B, iv, v). We used portions of an available 7680 member (20×384) dual unique indexed i5/i7 Nextera barcode library. However, barcoding oligo costs can be significantly reduced using a combinatorial indexing strategy.^28^

Tn5-based library preparation workflows (*e*.*g*. for single-cell libraries) often involve a bead-based cleanup and enrichment step of DNA templates prior to quantification, normalization, and tagmentation. This cleanup step is required to remove residual reagents and buffer components from dilute cDNA libraries^19^ but adds significant time and cost. We reasoned that the concentrated mutant amplicons (Figure 4A) used in our workflow could be diluted to reduce residual PCR components to avoid this step for uPIC–M, while still affording adequate amounts of DNA templates for tagmentation (typically performed at a template concentration of ∼0.1–1 ng/μL). Initial tests confirmed that for templates at identical concentrations, the yield after tagmentation and subsequent library amplification was comparable for purified and diluted samples (Figure S6A) and that 0.1–0.5 ng/μL template prior to tagmentation resulted in quantifiable libraries with similar size distributions (Figure S6B). We diluted all SpAP sublibrary plates 1:100 in H_2_O prior to tagmentation, instead of performing the time-consuming normalization of individual wells, yielding final concentrations of ∼0.1–0.5 ng/µL prior to tagmentation (for stock concentrations, see Table S6). In the case of greater variation (>5-fold) in sample well DNA concentrations, multiple dilutions of the same plate could easily be performed and processed for sequencing with only modest increases in time and cost.

### Automated plate processing

The above steps are labor-intensive if performed manually, but automated liquid handling techniques that are now used widely for single-cell sequencing applications can readily process >20×384 sample plates (or 7680 clones) for sequencing per day (see Materials and Methods). This approach allowed tagmentation and barcoding amplification of the 13×384 plate SpAP library in less than one day.^19^

### Sequencing library QC

The final step prior to pooled NGS involves evaluating library quality by quantifying the final concentration and distribution of fragment sizes. We pooled barcoded and amplified single clone libraries from each plate (i.e., one pooled sample per plate), purified and enriched them via a magnetic bead cleanup with size selection (see Materials and Methods), and then estimated fragment sizes and concentrations by microelectrophoresis (Figure 5). The library quality and concentration varied by sample plate (Figure S7, Table S6); 7/13 sublibraries contained clear fragment peaks at 400–500 bp and all samples contained measurable fragments between 400– 1000 bp without detectable contamination from low molecular weight sequencing adapters. This library quality allowed recovery of 65% of barcodes at high read depth across sublibrary plates (see below).

**Figure 5.**
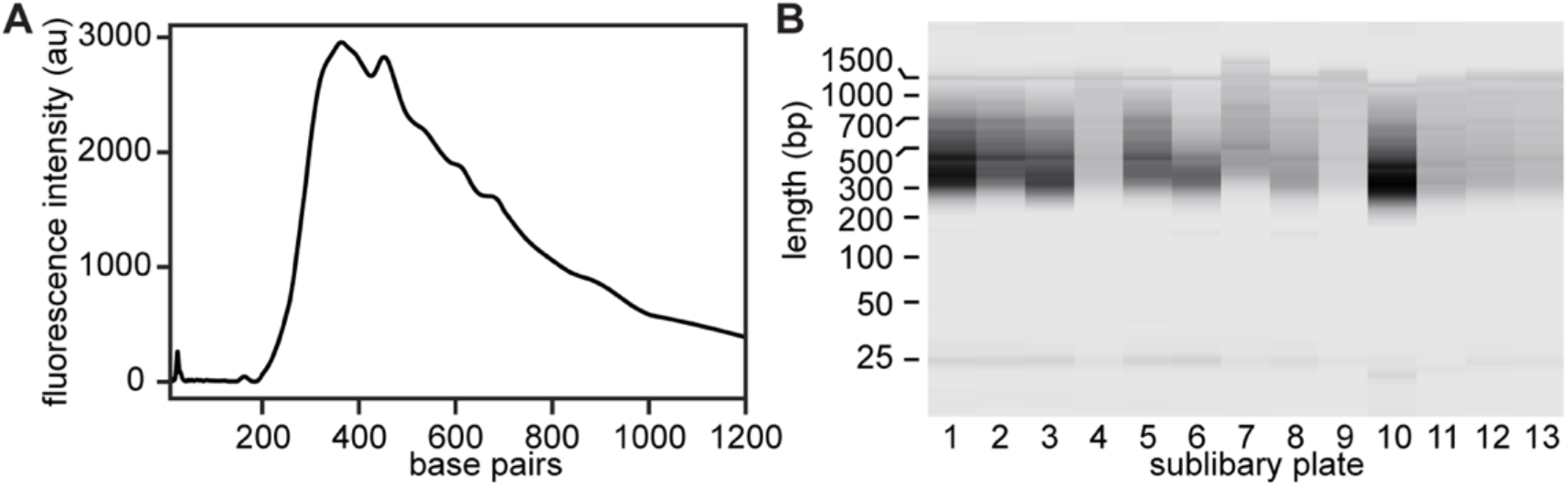
Sequencing library quality control results. **(A)** Plot of fluorescence (arbitrary units) vs. fragment length for sublibary 1 following tagmentation and barcoding amplification. See Figure S7 for analogous data for the other sublibraries. (**B**) Electropherograms of sublibraries 1–13 (see Table S6 for integrated peak concentrations).

### Sequencing library analysis

Analyzing results from NGS sequencing requires: (1) grouping reads by barcode, (2) eliminating barcodes with low coverage, (3) removing poor quality bases and residual adapter sequences from reads, (4) aligning reads to the “reference” ORF (here, the SpAP-eGFP amplicon sequence), and (5) identifying and evaluating sequence variants (Figure 6A, B).

**Figure 6.**
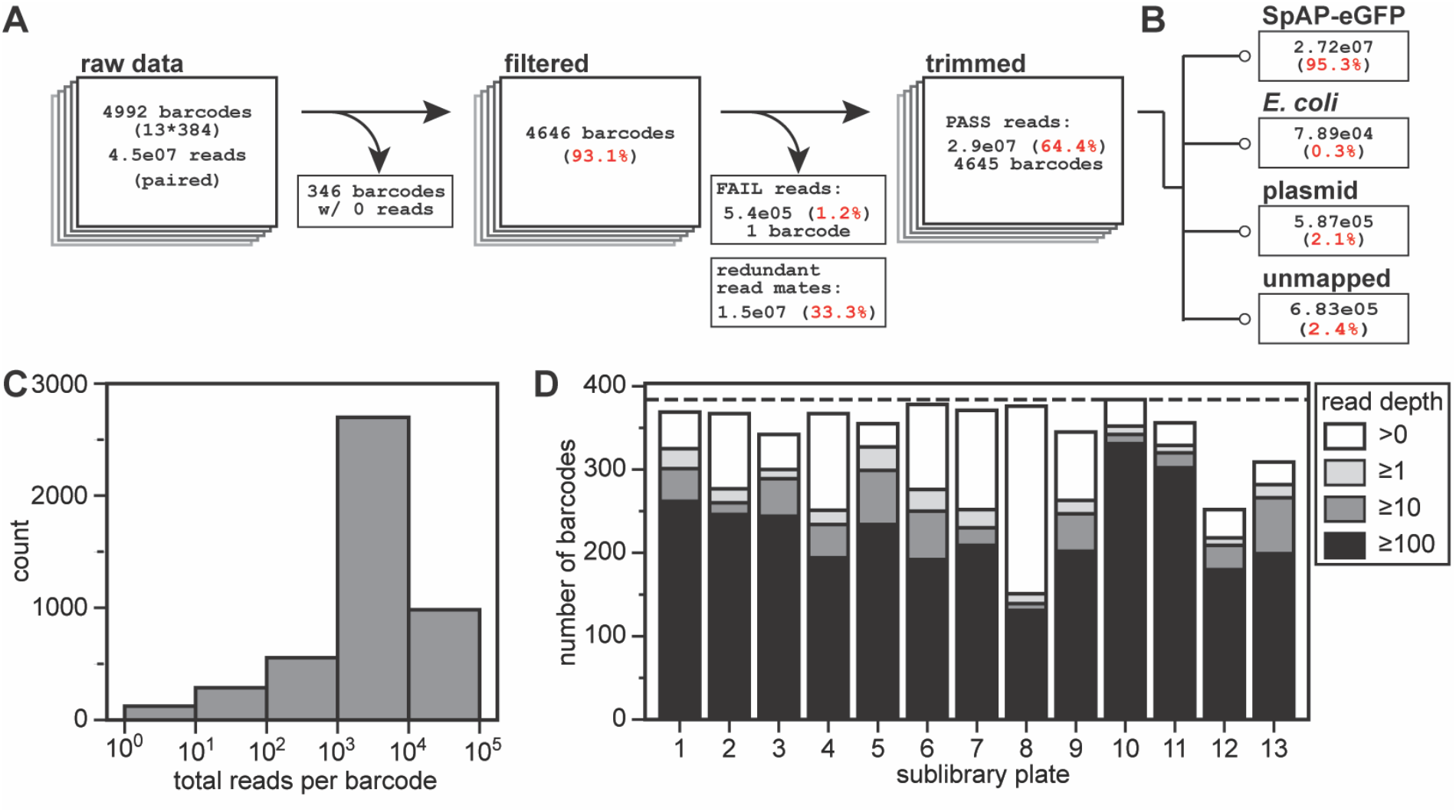
NGS data processing and read mapping pipeline and results for the SpAP scanning library. (**A**) Data processing steps and observed statistics. Raw FASTQ files (demultiplexed and unpaired) are filtered for barcodes containing 1 or more reads followed by adapter sequence trimming and pairing with read mates (if both reads are present and meet length/quality thresholds). Sequence-redundant readthrough read pairs are flagged at this stage and redundant read mates are discarded. (**B**) Trimmed and paired reads are mapped to the SpAP-eGFP amplicon, *E. coli*, and full plasmid genomes. (**C**) Histogram of total reads per barcode across all sublibraries following read trimming and pairing (n = 4645). (**D**) Barcode counts for each sublibrary plate at several read depth thresholds for the SpAP-eGFP ORF (>0 represents barcodes containing any mapped reads and remaining thresholds represent the minimum number of mapped reads at all positions; only barcodes containing at least one mapped read are included). The horizontal dashed line at 384 barcodes represents the maximum possible number of barcodes.

From a 25 million capacity MiSeq v3 (2×300 bp) run, we obtained 4.5×10^7^ total reads (read 1 and read 2) that were demultiplexed by the instrument using supplied i7/i5 barcodes (4992 barcodes total). We then discarded FASTQ files for barcodes with 0 reads (4646 retained barcodes, Figure 6A), trimmed off the universal Illumina adapter sequences, filtered reads based on quality scores and length using standard criteria,^29^ and paired reads with their mate if present. If paired reads were fully redundant, i.e. with readthrough to an adapter sequence on the opposing terminus, one mate was discarded (typically R2).^29^ We recovered 2.9×10^7^ reads after the trimming and associated filtering step (Figure 6A), with a median of 6×10^3^ reads per barcode (Figure 6C, Table 1). Even for sublibraries with relatively poor tagmentation yields (4, 7, 9, 11–13; Table S6), median reads per barcode were comparable (within <3-fold) to efficiently tagmented sublibraries (Table 1). This consistency was likely aided by sample normalization prior to sequencing, which accounted for differences in library concentrations.

**Table 1.**
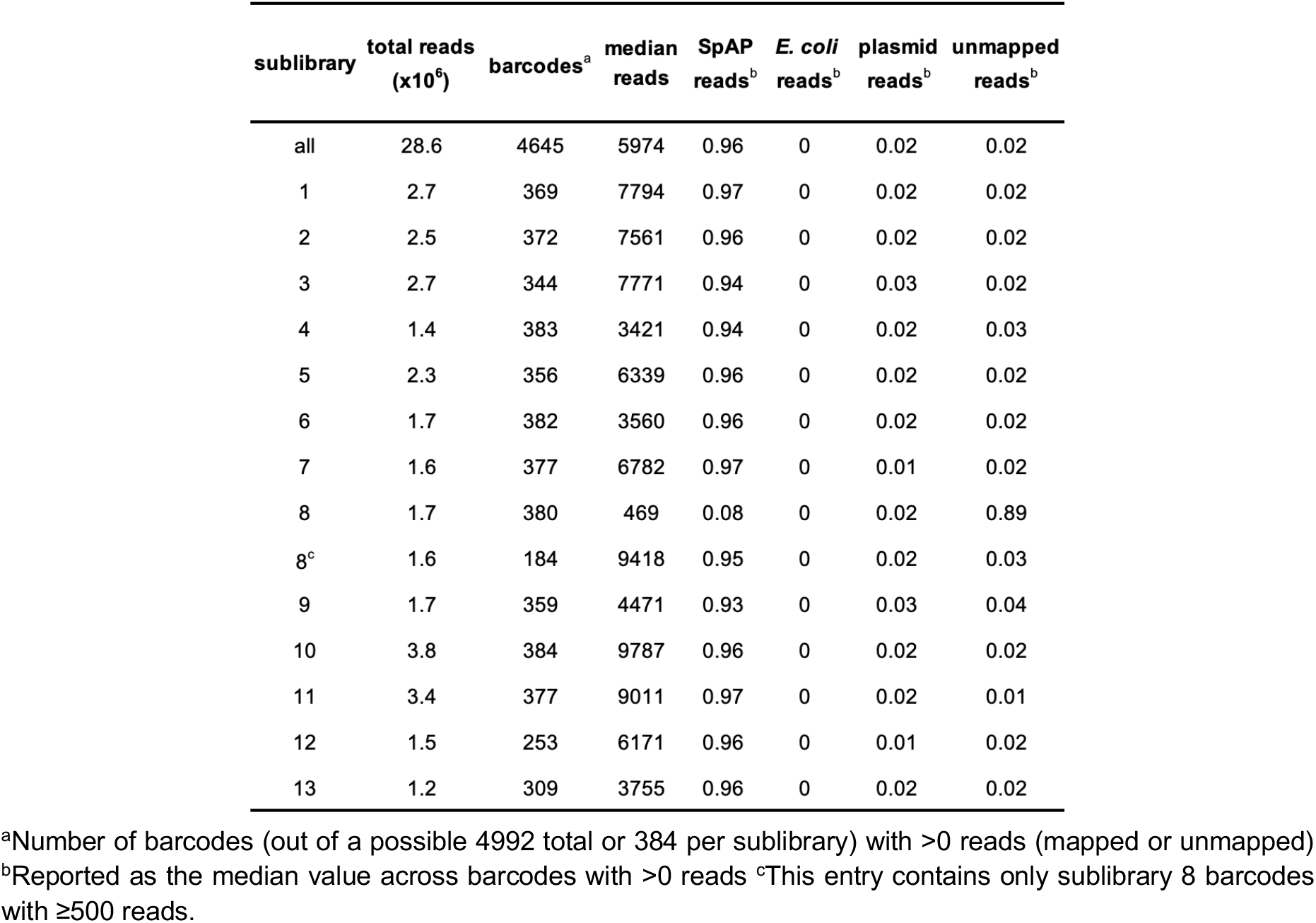
SpAP mutational sublibrary sequencing statistics.

Next, we mapped the reads to multiple reference genomes using the Burrows-Wheeler Aligner.^30^ For the SpAP mutagenesis library presented here, 95.3% of these reads mapped to the SpAP amplicon reference sequence (this sequence includes the 5’-UTR, eGFP fusion, and the 3’-UTR) and an additional 0.3% mapped to the *E. coli* genome. The remaining reads mapped to plasmid-derived sequences outside of the amplicon region (2.1%) or were unmapped (2.4%), likely representing either low quality reads or contamination from human or other sources (Figure 6B). The ratio of SpAP-eGFP to *E. coli* reads was highly consistent across sublibrary plates (Table 2). The read depth per barcode (calculated as the median read depth for all nucleotide positions of the SpAP-eGFP ORF *within* each barcode) varied from 0-1700 reads, with a median of 433 reads. Across the entire library, 65% of recovered barcodes were sequenced to a depth of ≥100 reads at all positions (Figure 6D, Table 1).

**Table 2.** SpAP read depth statistics.

| sublibrary | barcodes<sup>a</sup> | depth<br>≥1<sup>b</sup> | depth<br>≥10<sup>b</sup> | depth<br>≥100<sup>b</sup> | depth<br>≥1000<sup>b</sup> | kept<sup>c</sup> |
| --- | --- | --- | --- | --- | --- | --- |
| all | 4571 | 3603 | 3386 | 2926 | 0 | 3530 |
| 1 | 369 | 325 | 301 | 262 | 0 | 318 |
| 2 | 367 | 277 | 260 | 246 | 0 | 274 |
| 3 | 342 | 300 | 289 | 244 | 0 | 298 |
| 4 | 367 | 251 | 234 | 194 | 0 | 247 |
| 5 | 355 | 327 | 299 | 234 | 0 | 315 |
| 6 | 378 | 276 | 250 | 192 | 0 | 264 |
| 7 | 371 | 252 | 230 | 209 | 0 | 241 |
| 8 | 376 | 151 | 139 | 131 | 0 | 146 |
| 9 | 345 | 263 | 247 | 202 | 0 | 260 |
| 10 | 384 | 352 | 342 | 331 | 0 | 352 |
| 11 | 356 | 329 | 320 | 302 | 0 | 329 |
| 12 | 252 | 218 | 209 | 180 | 0 | 216 |
| 13 | 309 | 282 | 266 | 199 | 0 | 270 |

<sup>a</sup>Number of barcodes (out of a possible 4992 total or 384 per sublibrary) with ≥1 SpAP reads <sup>b</sup>Number of barcodes with at least this read depth at all positions in the SpAP-eGFP genome <sup>c</sup>Number of barcodes meeting the depth threshold of at least 1 read at all positions, and ≥10 reads at ≥95 % of positions. Only barcodes meeting this depth threshold were carried forward for subsequent variant analyses.

### Quantifying the yield of single mutants

The next stage of assembling a ready-to-use library for high-throughput biochemistry assays is to identify the clones containing single mutations and map each mutant to plate-well locations. We selected barcodes meeting the following depth threshold for further analysis: ≥1 read at all genome positions *and* ≥10 reads at ≥95% of all reference genome positions. Of 4645 barcodes with ≥1 mapped read, 3530 (76%) met this threshold across the entire library. To detect variants associated with each barcode in batch, we applied a SAMtools module^31,32^ to process all mapped reads and generate output files (*variant call files*, .vcf) for each barcode containing SpAP-eGFP variants (single nucleotide substitutions, indels, or null if WT). WT and indel-containing barcodes were discarded (Figure 7A). For barcodes containing single nucleotide substitutions, we determined the corresponding codon and amino acid changes, assessed whether observed substitutions were *intended* (correct mutant identity and sublibrary) or *unintended*, and evaluated whether these barcodes contained single, double, or triple and greater numbers of amino acid substitutions. We also stored variant quality statistics at this stage, including the number of forward and reverse reads containing variant *vs*. WT nucleotide sequence. Among barcodes containing single mutants, most observed mutations were intended (97%) (Figure 7A).

**Figure 7.**
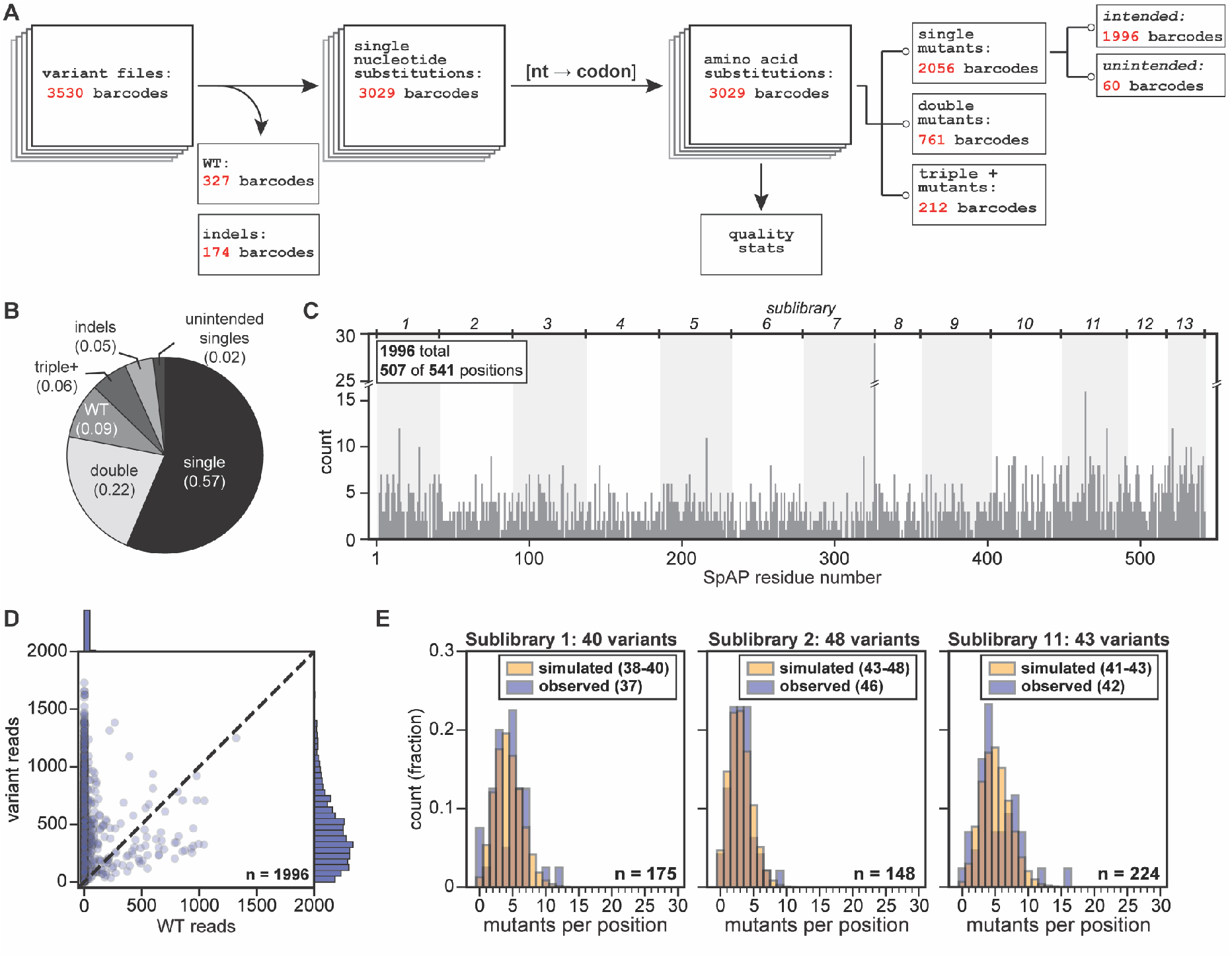
Characterization of the SpAP alkaline phosphatase scanning mutant library created with uPIC–M. (**A**) Overview of variant detection analyses and calculated yields (red) for the SpAP mutant library. (**B**) Overall distribution of single mutants, WT, double mutants, triple and greater mutants, and indels across all mutational sublibraries (indel count reflects variants containing one or more indels). (**C**) Location and frequency of intended single mutants across the entire SpAP-eGFP ORF. (**D**) Scatter plot and histograms of variant reads vs. WT reads for all intended single mutants. (**E**) Comparison of simulated and observed single mutant frequency distributions for three sublibraries. Legend specifies the observed yield of unique single mutants and simulated 95% confidence interval from 1000 events; ‘n’ indicates the total number of observed intended single mutants. Results for all sublibraries are shown in Figure S9.

Across all sublibraries, single mutants comprised 57% of the clones, ranging from 52–68% within each sublibrary (Figure 7B; Table 3), similar to the results of small-scale testing (60%, Table S4) and within the range of mutant picking simulations (10–100%; see “Simulated mutant sampling to predict screening requirements”) (Table 3). Double, and triple and greater mutants comprised 28% of sequenced clones (Figure 7B), higher than the 5% observed during small-scale testing. This higher percentage may arise in part from cross-contamination between single mutant clones during plate handling steps prior to barcode introduction, which was absent during small scale testing. Variant:WT read ratios across single, double, and triple and greater mutants are consistent with this model (Figure S8).

**Table 3.** Variant content of the SpAP scanning library.

| sublibrary | residues | total positions | barcodes<sup>a</sup> | single total | single intended | single unintended | double | triple+ | indels | WT |
| --- | --- | --- | --- | --- | --- | --- | --- | --- | --- | --- |
| all | 2–542 | 541 | 3530 | 2056 | 1996 | 60 | 761 | 212 | 174 | 327 |
| 1 | 2–41 | 40 | 318 | 178 | 175 | 3 | 61 | 18 | 20 | 41 |
| 2 | 42–89 | 48 | 274 | 154 | 148 | 6 | 66 | 17 | 8 | 29 |
| 3 | 90–137 | 48 | 298 | 168 | 161 | 7 | 65 | 20 | 14 | 31 |
| 4 | 138–185 | 48 | 247 | 135 | 131 | 4 | 43 | 27 | 24 | 18 |
| 5 | 186–232 | 47 | 315 | 175 | 169 | 6 | 85 | 11 | 21 | 23 |
| 6 | 233–279 | 47 | 264 | 141 | 138 | 3 | 60 | 23 | 23 | 17 |
| 7 | 280–326 | 47 | 241 | 141 | 134 | 7 | 59 | 12 | 5 | 24 |
| 8 | 327–356 | 30 | 146 | 94 | 91 | 3 | 21 | 5 | 2 | 24 |
| 9 | 357–402 | 46 | 260 | 148 | 142 | 6 | 50 | 11 | 16 | 35 |
| 10 | 403–448 | 46 | 352 | 210 | 204 | 6 | 81 | 24 | 7 | 30 |
| 11 | 449–491 | 43 | 329 | 230 | 224 | 6 | 60 | 15 | 5 | 19 |
| 12 | 492–517 | 26 | 216 | 120 | 120 | 0 | 52 | 20 | 12 | 12 |
| 13 | 518–542 | 25 | 270 | 162 | 159 | 3 | 58 | 9 | 17 | 24 |

<sup>a</sup>Number of barcodes meeting the depth threshold of at least 1 read at all positions, and $\geq 10$ reads at $\geq 95$ % of positions.

Next, we examined the identity and location of single mutant variants and found that mutants were evenly distributed with minimal positional bias (Figure 7C). Overall, we recovered 507 of 541 desired single mutants (94%), with coverage ranging from 87–98% within sublibraries (Table S7). These data demonstrate that uPIC–M is capable of producing user-defined single mutant libraries at high coverage.

### Assessment of single mutant purity

High-throughput biochemistry demands different levels of mutant purity depending on the application. Quantitative measurements of variant stabilities or ligand affinities can tolerate low amounts of contamination, as this contamination leads to accordingly low errors on thermodynamic parameters. By contrast, measurements of enzyme turnover are highly sensitive to contamination, as a small (∼1%) fraction of WT enzyme could dominate apparent rates when attempting to measure a catalytically-impaired mutant with activity that is <<1% of WT.

Above, we classified all barcodes containing only one amino acid mutation as single mutants without considering the fraction of mutant reads at that position. Here, we assess mutant purity by quantifying the number of forward and reverse reads containing either the mutant or WT nucleotide at the mutated position. Across this library, single mutants contained a wide range (10–1000) of variant reads, and relatively few but detectable WT reads (Figure 7D). To calculate mutant purities, we devised a quality threshold representing the minimum number of variant reads *and* the minimum ratio of variant:WT reads (Table S7); as many barcodes contained 0 WT reads, this threshold represents a lower limit on single mutant purities. Library yield dropped from 507 mutants (94%) to 498 (92%) mutants and 484 (89%) mutants when applying thresholds of 10 and 100, respectively.

### Evaluation of method performance

To measure the success of uPIC–M compared to picking simulations, we calculated expected unique mutant yields per SpAP sublibrary. We repeated simulations, this time substituting values for experimental variables that were assumed using only generic estimates in the initial simulations described above. These parameters were: 1) the number of sequenced barcodes per sublibrary, which is then used as the number of simulated draws, and for each sublibrary was fewer than the 384 possible, 2) observed single mutant frequency, as this defines the chance of drawing a single mutant among a pool containing multiple categories of variants, and 3) the total number of possible unique mutants, which varied by sublibrary (Table S1) and is proportional to the number of draws necessary to achieve a desired level of library coverage.

For each sublibrary, we ran 10^4^ picking simulations to estimate the expected median number of unique mutants (and 95% confidence interval) and distribution of mutants per position (Figure 7E, selection of 3 sublibraries; Figure S9, all sublibraries). Observed *unique* single mutant yields matched expectations for 10 of 13 sublibraries (2–8, 11–13) (Table S8). The remaining 3 plates (1, 9, 10) yielded one fewer single mutant than expected at the 95% confidence interval, suggesting that an unequal abundance of some single mutants slightly decreased library coverage. Together, these results establish that the picking simulations presented here can accurately guide experimental pipelines for generating uPIC–M libraries.

## CONCLUSIONS

uPIC–M delivers user-designed, clonal, single mutant libraries at a significant savings of time and cost compared to conventional mutagenesis by combining commercially available oligonucleotide arrays with commonly-used automated liquid handling platforms and barcoded Illumina sequencing strategies. Here, we used this method to rapidly generate a single mutant scanning library of a 541 amino acid enzyme and achieved a >90% yield of desired variants. This method is immediately applicable to new protein targets, and we provide a detailed workflow for rigorous characterization of library sequences using a collection of open-source and custom data analysis tools.

In future work, uPIC–M can readily be extended to a variety of applications beyond generating single mutant libraries. The relatively long mutagenic primers used in QuikChange-HT mutagenesis hybridize efficiently and specifically even in the presence of several nucleotide substitutions, making it possible to introduce multiple mutations within the same mutagenic window.^33,34^ uPIC–M’s window design strategy can also be adapted to create deletion or insertion libraries using assembly-based strategies in pool.^35,36^ A software tool under development by the Fordyce group will facilitate the design of single mutant and other variant libraries.^37^ Finally, uPIC–M can be readily adapted for sequencing with other Illumina instruments or long-read sequencing approaches.^38^

High-throughput biochemistry reveals biophysical insights on an unprecedented scale, but constructing variant libraries has become the new rate-limiting step.^5^ The ability of uPIC–M to generate the required variant libraries rapidly and efficiently will expand HTB to the study of new protein targets, systems, and questions.

## MATERIALS AND METHODS

### Description of plasmid

The plasmid mutagenized here encoded the SpAP alkaline phosphatase family monoesterase^39,40^ (Uniprot KB – A1YYW7) fused to a C-terminal eGFP *via* a 10 amino acid ser-gly linker. SpAP residues 1–540 (full-length *sans* signal peptide, original numbering with signal sequence = 20–559) were subcloned into the manufacturer-supplied plasmid from the PURExpress® In Vitro Protein Synthesis Kit (New England Biolabs, Ipswich, MA, USA) using the Gibson assembly method with synthetic *E. coli* codon-optimized SpAP DNA (IDT, Coralville, IA, USA). Full plasmid map (Figure S10), DNA sequence (Figure S11), and the SpAP-eGFP fusion protein sequence (Figure S12) are provided in supplemental material.

### Design of mutagenic oligo arrays

Oligo arrays were designed using the Agilent Technologies (Santa Clara, CA, USA) eArray web program (https://earray.chem.agilent.com/earray/). The full sequence of the PURExpress-SpAP-eGFP was provided as input to the eArray software (Figure S11), and mutational regions were manually adjusted until primer sequences for all sublibrary mutational windows (13 total) passed thermodynamic thresholds calculated by this software. Oligos were selected to mutate all non-valine residues from positions 2–326 to valine; all valine residues from positions 2–326 to alanine; all non-alanine residues from positions 327–542 to alanine; and all alanine residues from positions 327–542 to synonymous alanine codons, corresponding to 541 total mutants (Table S1). Each mutagenic oligo was synthesized in duplicate within a 7500 oligo capacity high-fidelity array (Agilent Technologies, see Table S9 for array sizes and sample pricing). The forward and reverse primer sequences (Table S1) required to amplify each mutational window from the pooled array were also obtained from the eArray design output and were purchased from IDT. The full array sequence is provided in an accompanying data repository (https://osf.io/k3rjy/).

### Preparation of sublibrary mutagenic primer pools and PCR mutagenesis

Lyophilized oligo arrays were resuspended in 200 μL 10 mM Tris-HCl, pH 8.5 (EB), and then further diluted 1:100 with EB. Sublibrary oligo pools were amplified individually using window-specific primer pairs and KAPA HiFi HotStart ReadyMix (Roche, Indianapolis, IN, USA) with a final template concentration of 1:2000 resuspended oligo array and annealing temperatures of either 60 °C or 65 °C (depending on performance of individual primer pairs) for 25 PCR cycles. Resulting PCR products (“sublibrary mutagenic primers”) were purified using the StrataPrep PCR Purification Kit (Agilent Technologies) and analyzed for quality and concentration using TapeStation electrophoresis with HSD1000 ScreenTapes (Agilent Technologies). For initial mutagenesis reactions, primer stocks were normalized to a uniform concentration, measured by UV absorbance, of 15 nM, that of the lowest concentration sublibrary pool (Table S2). PCR mutagenesis was performed using the QuikChange Lightning enzyme (Agilent Technologies) at an annealing temperature of 60 °C for 18 cycles with the following components: 2.5 μL 10X manufactuer-supplied buffer, 1 μL supplied dNTP mix, 0.75 μL QuikSolution additive, 1 μL of 25 ng/μL PURExpress-SpAP-eGFP template plasmid, 15 μL 14.7 nM sublibrary pool, 1 μL QuikChange Lightning enzyme, 3.75 μL H_2_O. Following PCR, template WT plasmid was digested by the addition of 1 μL DpnI (Agilent Technologies) for 5 min at 37 °C. For sublibraries that provided an insufficient number of transformants, mutagenesis was repeated using the undiluted purified sublibrary primer pools.

### Transformation, plating, colony picking & growth

DpnI-digested PCR mutagenesis reactions were transformed into chemically-competent NEB 5-alpha *E. coli*. Transformations were plated on 15 cm LB agar plates containing 100 µg/mL ampicillin and grown overnight at 37 °C. Transformation ratios of 1:20–1:3.33 PCR product:cells were used to obtain a desired yield of ∼400–500 colonies per 15 cm plate. Colonies were picked manually (sublibraries 1 and 5 only) or using a PIXL robotic colony picker (Singer Instrument Company, Somerset, UK) at the Stanford University School of Medicine Genome Technology Center (Palo Alto, CA, USA). Single colonies were picked from source LB agar plates into 384 well (120 μL) destination microwell plates containing 60 μL LB containing ampicillin. At least 384 colonies were picked and grown from each sublibrary window (Figure 2). Microwell plates were sealed with gas-permeable AeraSeal film (MilliporeSigma, Burlington, MA, USA) and grown to saturation with shaking at 37 °C.

### qPCR detection of E. coli genomic DNA

*E. coli* cultures from clonal mutants were pooled and diluted from 10^1^ to 10^4^-fold to assay for genomic DNA concentration using qPCR with the commercial NEB Luna 2X MasterMix. A previously reported primer set to the *rodA* gene was used (Forward: 5’-GCAAACCACCTTTGGTCG-3’; Reverse: 5’-CTGTGGGTGTGGATTGACAT-3’).^41^ Library samples were quantified using a standard curve of purified *E. coli* O157:H7 genomic DNA (Zeptometrix, Buffalo, NY, USA) at concentrations of 0.0001–1 ng/μL.

### Preparation of enzyme ORF amplicons

Saturated clonal *E. coli* cultures in 384 well plates were diluted 1:1000 with H_2_O, by serial dilution using a 96-well Rainin Liquidator (Mettler-Toledo, Columbus, OH, USA). Dilutions and additional amplicon preparation steps were performed in 384 well Bio-Rad HSP3801 plates (Bio-Rad, Hercules, CA, USA). Primers were designed to amplify a 2525 bp region including the SpAP-eGFP ORF and 5’- and 3’-UTR segments (Figure S10). Forward (5’-gatctacactctttccctacacgacgctcttccgatctCCCGCGAAATTAATACGACTCACTATAGG-3’) and reverse (5’-gtctcgtgggctcggagatgtgtataagagacagGCACCACCTTAATTAAAGGCCTCC-3’) primers also contained Illumina Read 1 and Read 2 overhangs, respectively, shown in lowercase. Diluted cultures were used as PCR templates and amplified with KAPA HiFi HotStart polymerase at a scale of 4 μL: 0.8 μL 1:1000 dilute culture template, 2 μL KAPA HiFi HotStart ReadyMix (Roche), 0.96 μL H_2_O, 0.12 μL 10 μM forward primer, 0.12 μL 10 μM reverse primer (see sequences above). Thermal cycling conditions were as follows: 95 °C, 5 min; 25x[98 °C, 20 s; 60 °C, 15 s; 72 °C, 2 min]; 72 °C, 2 min.

### Fluorescence quantification of amplicon DNA

Amplicon DNA concentrations after PCR were determined using the Quant-iT PicoGreen dsDNA assay (Thermo-Fisher Scientific, Waltham, MA, USA). In brief, a master mix containing 1:200 PicoGreen reagent (from stock concentration as supplied) to 10 mM Tris-HCl, pH 8.5, was prepared immediately before use and kept from light. Amplicon samples were prepared by the addition of 1.5 μL of 1:5 PCR reaction:H_2_O to 34 μL master mix. Standards were prepared by the addition of 1.5 μL λ phage DNA ranging in concentration from 0–100 ng/μL to 34 μL master mix. Standard curves were included, in duplicate, on each sample plate. Samples were incubated ∼5 min at room temperature, and read by fluorescence (λ_ex_ = 480 nm, λ_em_ = 520 nm) using a Synergy H1 plate reader (BioTek, Winooski, VT, USA).

### Tn5 tagmentation

A library preparation procedure adapted from Picelli et. al.,^19,26^ with modifications, was used to generate barcoded, sequencing-ready libraries. Commercial pA-Tn5 (protein A-Tn5) was purchased pre-loaded with sequence adapters from Diagenode (Denville, NJ, USA), and diluted to a working concentration of 1:50 with dilution buffer (40 mM Tris-HCl, pH 7.5, 40 mM MgCl_2_). Amplicons in 384 well plates were diluted 1:100 in H_2_O to provide template concentrations suitable for tagmentation. For Tn5 reactions, 1.2 μL of master mix (0.2 μL 1:50 pA-Tn5; 1 μL 1.6x buffer containing 16 mM Tris-HCl, pH 8.0, 8 mM MgCl_2_, and 16% (v/v) dimethylformamide, sparged with N_2_ and added immediately prior to use) was added to each well of a new 384 well plate using a Mantis microfluidics liquid handler robot (Formulatrix, Bedford, MA, USA). Amplicon templates were added to Tn5 reaction mixtures (0.4 μL 1:100 template each) using a Mosquito LV pipetting robot (SPT Labtech, Boston, MA, USA). Reaction plates were sealed, briefly vortexed, collected by centrifugation, and incubated for 7 min at 55 °C. Tn5 reactions were stopped by the addition of 0.4 μL of 0.1% (w/v) sodium dodecyl sulfate, using the Mantis liquid handler.

### i7/i5 barcoding PCR and library cleanup

Tagmented libraries were barcoded and amplified using KAPA HiFI polymerase and a collection of Nextera XT 12mer dual unique index sequencing primers (purchased from IDT and supplied by CZ-Biohub). First, 1.2 μL of a master mix containing 0.08 μL KAPA HiFi (1 U/μL), 0.5 μL 5X buffer (manufacturer supplied), 0.12 μL 10 mM dNTP mix (2.5 mM each), and 0.2 μL H_2_O was added to each 2 μL SDS-halted Tn5 reaction using the Mantis liquid handler. Next, 0.8 μL of unique i5/i7 primer mix (2.5 μM each) was transferred from source plates to sample mixture using the Mosquito instrument. Reaction plates were sealed, briefly vortexed, collected by centrifugation, and amplified with the following thermal cycler conditions: 72 °C, 3 min; 95 °C, 30 s; 12x[98 °C, 10 s; 55 °C, 15 s; 72 °C, 1 min]; 72 °C, 5 min. Resulting libraries were pooled and treated with AMPure XP magnetic beads at a ratio of 0.8:1 beads:sample volume to purify DNA (Beckman Coulter, Brea, CA, USA). Library yield and quality was determined by TapeStation electrophoresis with HSD1000 ScreenTapes (Agilent Technologies).

### Next-generation sequencing

Sequencing was performed by SeqMatic (Fremont, CA, USA). Libraries were sequenced using Miseq v3 2×300 bp, with the addition of 1% PhiX control DNA. Samples were submitted with i7/i5 barcodes corresponding to each tagmented mutant and demultiplexed by the instrument.

### Picking simulations

Pools of mutants were simulated as numeric lists containing unique elements equal in number to desired simulated mutant pool. The *random* module (Python 3^42^) was used to sample from this pool pseudo-randomly, thereby simulating a large (compared to sampling events) mutant pool with identical distributions of each unique mutant (https://github.com/FordyceLab/uPICM). Simulation of sampling from a variant pool containing additional non-single mutants was accomplished by the above strategy with the addition of a preceding step. This step introduced a pseudo-random draw from a pool containing specified fractions of single mutants (0.1–1.0) and non-single mutants (0–0.9), with only draws picking among the single mutant fraction carried forward.

### NGS data processing, and analysis

NGS data were processed using open source software tools, executed with the Snakemake workflow tool.^43^ First, sequences of Illumina adapters were trimmed and redundant (read-through) read mates were disposed from demultiplexed fastq read files using the Trimmomatic package.^29^ Trimmed reads were aligned to the PURExpress-SpAP-eGFP plasmid, and separately, the full *E. coli* genome (NCBI Reference Sequence: NC_000913.3) using BWA-MEM,^30^ with the output mapped, sorted, and indexed with SAMtools.^31^ Variant base calls were identified against the PURExpress-SpAP-eGFP plasmid genome using the BCFtools utility of the SAMtools package.^32^ Alignments were visualized using Integrative Genomics Viewer.^44^ Subsequent analyses were performed with custom code using Python 3, available at (https://github.com/FordyceLab/uPICM). Raw sequencing files are provided in our data repository (https://osf.io/k3rjy/).

## ACKNOWLEDGEMENTS

We thank Daniel Mokhtari for assistance with data analysis, Agilent Technologies for generously providing oligo arrays, Dr. Robert St. Onge and the Stanford Genome Technology Center for assistance with robotic colony picking, and members of the Herschlag and Fordyce research groups for critical discussions and reviewing the manuscript. This work was supported by the NIH grant R01 (GM064798) awarded to D.H. and P.M.F., an Ono Pharma Foundation Breakthrough Innovation Prize, and the Gordon and Betty Moore Foundation (grant number 8415). P.M.F. is a Chan Zuckerberg Biohub Investigator.

## FOR TABLE OF CONTENTS ONLY

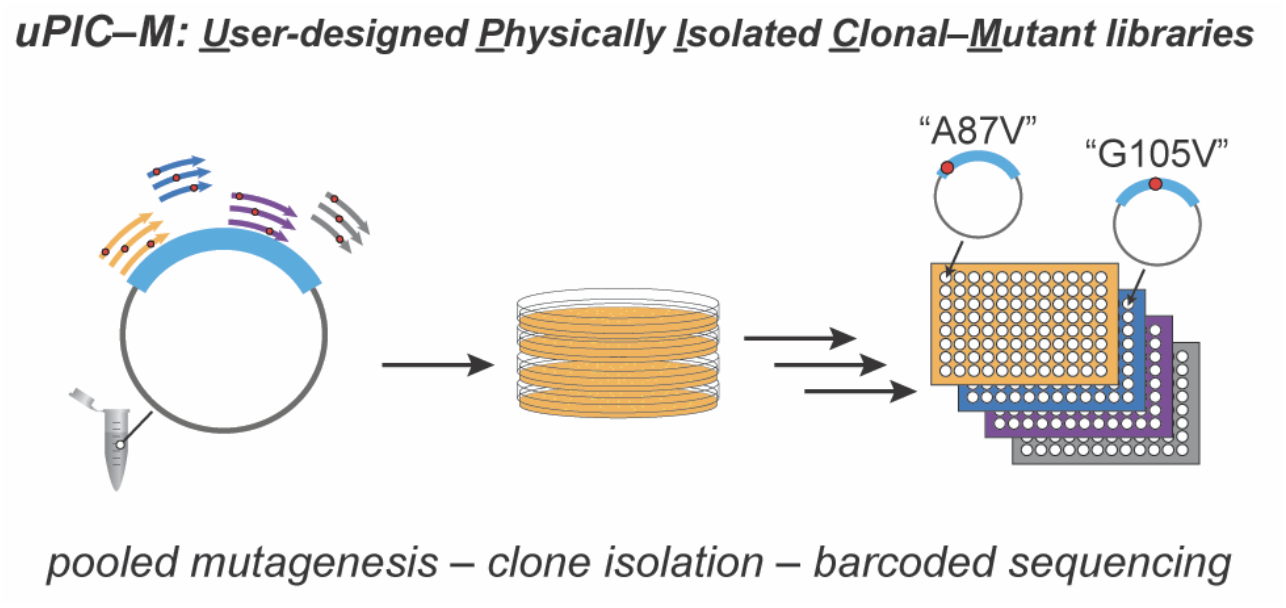

## Notes

https://osf.io/k3rjy/

### Competing Interest Statement

The authors have declared no competing interest.

