## Supporting Information for "uPIC–M: efficient and scalable preparation of clonal single mutant libraries for high-throughput protein biochemistry"

**TABLE OF CONTENTS**

**Figure S1**. Timeline of uPIC–M library generation. 3
**Figure S2**. Amplification of window-specific sublibrary pools from an oligo array. 4
**Figure S3**. Quantification of *E. coli* genomic DNA in diluted mutant culture templates. 5
**Figure S4**. Selection of PCR conditions for SpAP mutant amplicons. 6
**Figure S5**. Quantification of amplicon DNA concentrations per sublibrary plate. 7
**Figure S6**. Testing Tn5 tagmentation reaction conditions. 8
**Figure S7**. Electropherograms of tagmented and amplified mutant sublibraries. 9
**Figure S8**. Histogram of variant:WT read ratios among single, double, and triple and greater mutants. 10
**Figure S9**. Comparison of observed and simulated single mutant frequency distributions. 11
**Figure S10**. Plasmid map of PURExpress-SpAP-eGFP. 12
**Figure S11**. Complete DNA sequence of PURExpress-SpAP-eGFP plasmid. 13
**Figure S12**. Protein sequence of SpAP-(10mer linker)-eGFP. 14
**Table S1**. Oligo array and window design details for SpAP scanning mutant library. 15
**Table S2**. Concentration of purified sublibrary mutagenic primer pools. 16
**Table S3**. Expected mutant yields from simulations of mutant sampling. 17
**Table S4**. Variant composition of small-scale QuikChange-HT reactions. 18
**Table S5**. Sublibrary transformation and colony picking results. 19
**Table S6**. Amplicon DNA and library concentration statistics. 20
**Table S7**. Unique single mutant yields for the SpAP scanning library. 21
**Table S8**. Comparison of uPIC–M performance with simulated picking experiments, per sublibrary. 22
**Table S9**. Oligo array price summary. 23

**Figure S1**. Timeline of uPIC–M library generation.

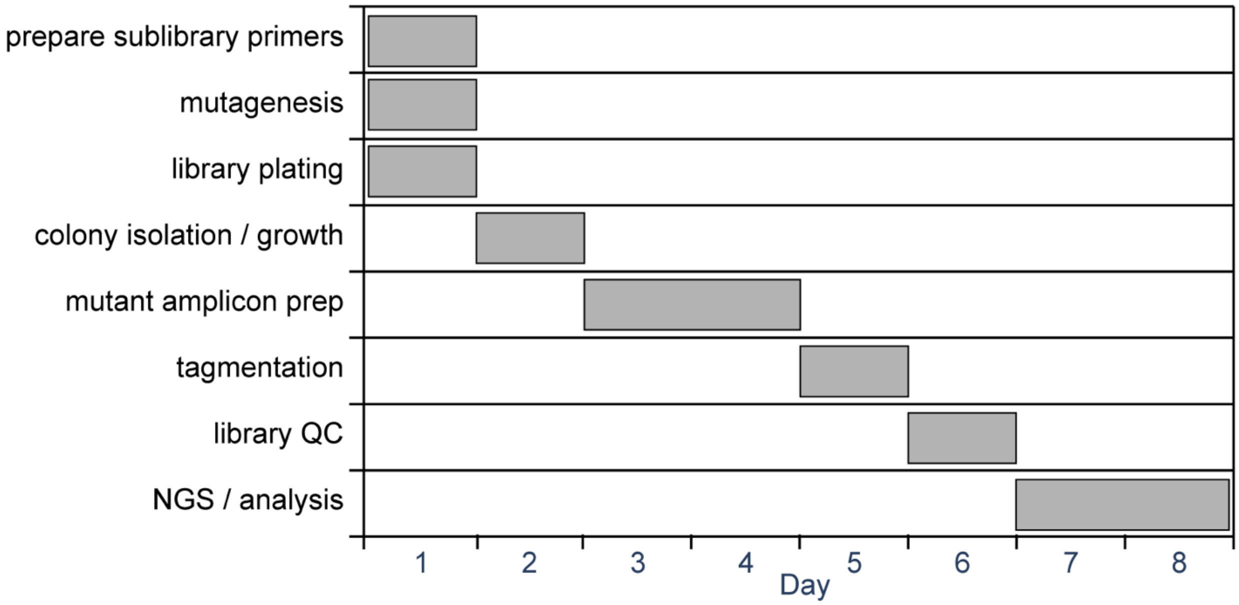

**Figure S2**. Amplification of window-specific sublibrary pools from an oligo array.
(**A**) Microelectrophoresis results for PCR-amplified sublibrary mutagenic primers after column purification. (**B**) Plot of predicted and observed lengths for mutagenic primer pools corresponding to sublibrary windows 1–13.

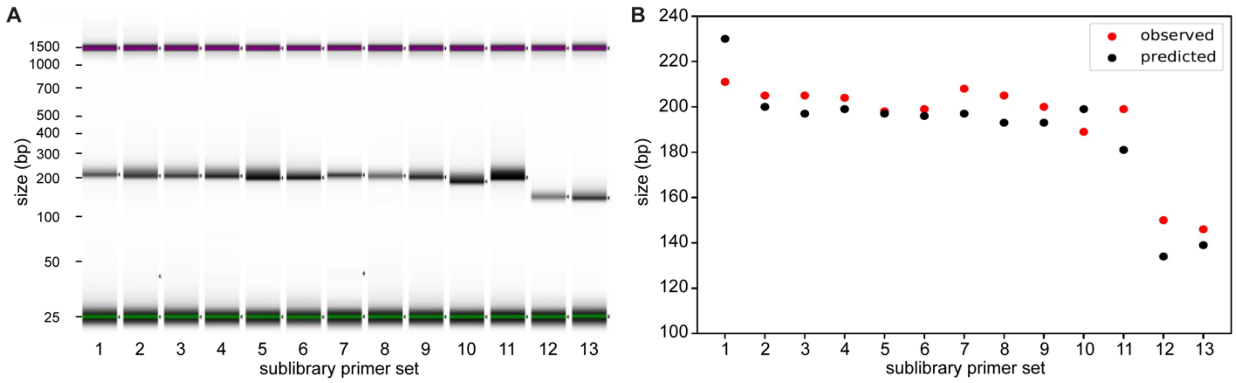

**Figure S3**. Quantification of *E. coli* genomic DNA in diluted mutant culture templates. (**A**) Standard curve of *E. coli* genomic DNA concentration measured by qPCR using previously reported primers to the *rodA* gene. Each point represents the average of 4 technical replicates. (**B**) Measurement of *E. coli* genomic DNA concentrations in diluted mutant cultures by qPCR. Six saturated mutant cultures were serially diluted in $H_2O$ and assayed alongside the standard curve in (**A**). The average of two technical replicates is plotted for each of the six biological replicates at each dilution. The black horizontal line represents the median across biological replicates.

**A**

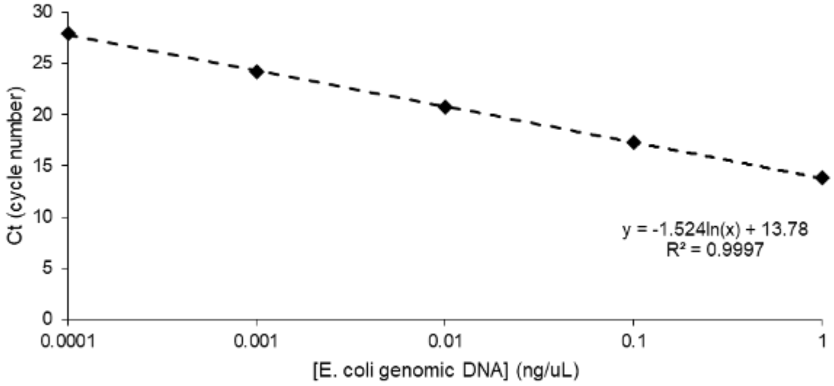


**B**

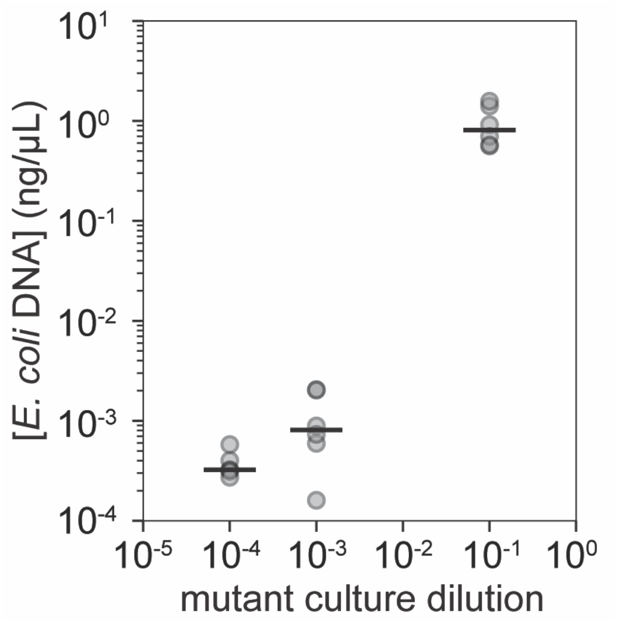

**Figure S4**. Selection of PCR conditions for SpAP mutant amplicons.
Twelve sample wells containing *E. coli* cultures of SpAP mutants grown to saturation were selected from sublibrary plates 1 and 5 (6 each). Cultures were then diluted from 1:10 to $1:10^4$ with $H_2O$ and used as PCR templates for 18 or 25 cycles of PCR amplification using KAPA HiFi polymerase. Primers were selective for a 1737 bp region of the PURExpress-SpAP-eGFP plasmid (F: 5'-CCCGCGAAATTAATACGACTCACTATAGG 3'; R: 5'- CTTGCTCACCATGCCACTG -3'). Following PCR, samples were diluted with $H_2O$ and loading buffer and run at equal volumes on a 0.8% TAE agarose gel.

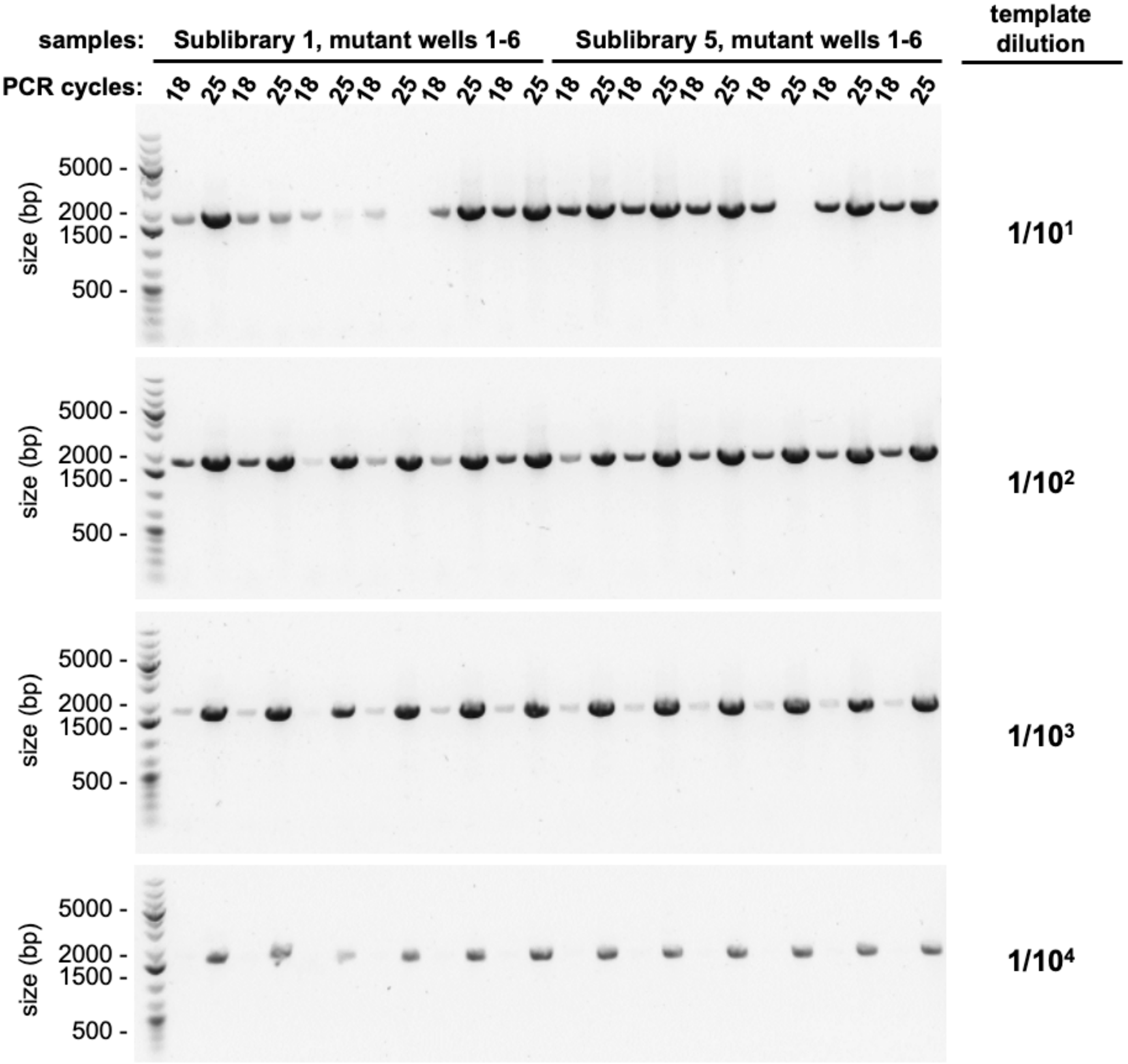

**Figure S5**. Quantification of amplicon DNA concentrations per sublibrary plate.
DNA concentrations were measured for each sublibrary plate (184 or 368 sample wells out of 384 possible) using the PicoGreen fluorescence assay. Amplicon samples were diluted 5-fold with $H_2O$ and measured alongside a λ phage DNA standard curve.

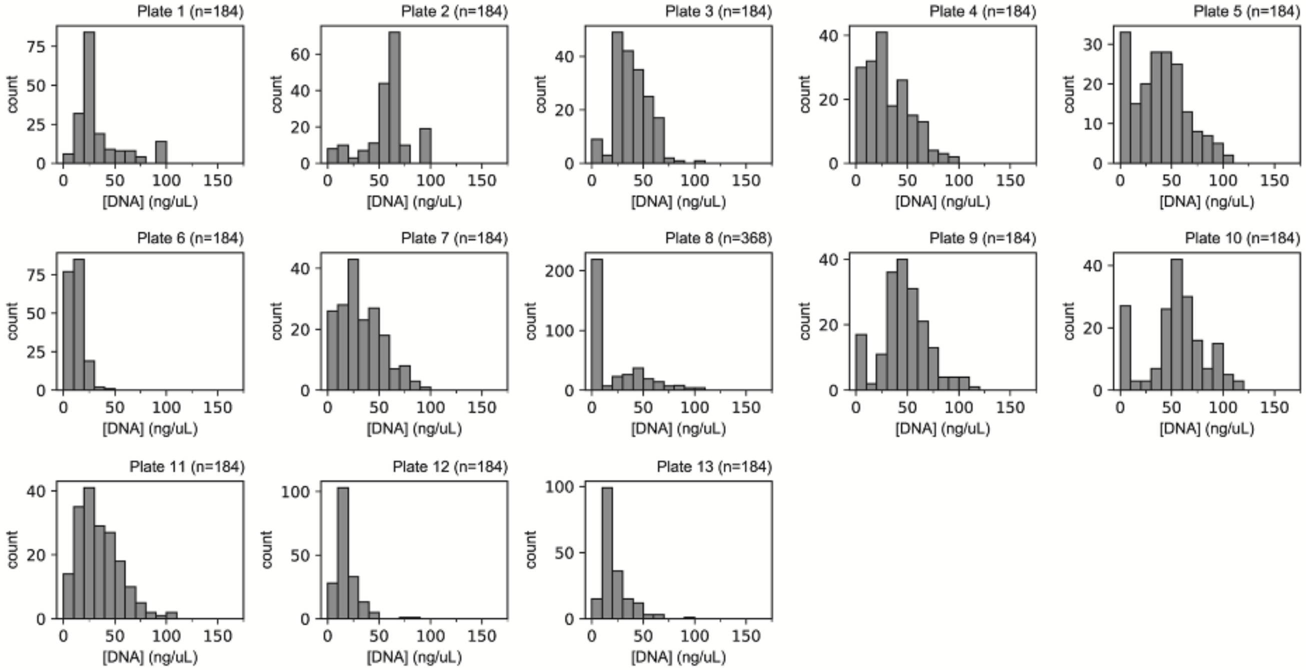

**Figure S6**. Testing Tn5 tagmentation reaction conditions.
(**A**) Comparison of library content and concentration with or without AmpureXP bead cleanup of Tn5 templates. Following mutant amplicon preparation, two unique scanning library sample plates were either purified by AmpureXP bead cleanup or simply diluted prior to Tn5 tagmentation. The Tn5 stock used here was in-house purified from the protocol of Picelli et. al., 2014. (**B**) Comparison of library content and concentration at template DNA concentrations of 0.1–0.5 ng/uL, Tn5 enzyme dilutions of 1/50 and 1/100, and tagmentation times of 3 and 7 minutes. The Tn5 template was purified SpAP amplicon DNA. Samples were allowed to react for specified times, quenched, and then amplified by library preparation PCR (see Materials and Methods). Amplified libraries were purified by AmpureXP bead cleanup and analyzed by microelectrophoresis. In both (**A**) and (**B**), DNA peak concentrations represent the integrated signal of the entire library peak (~200–1000 bp).

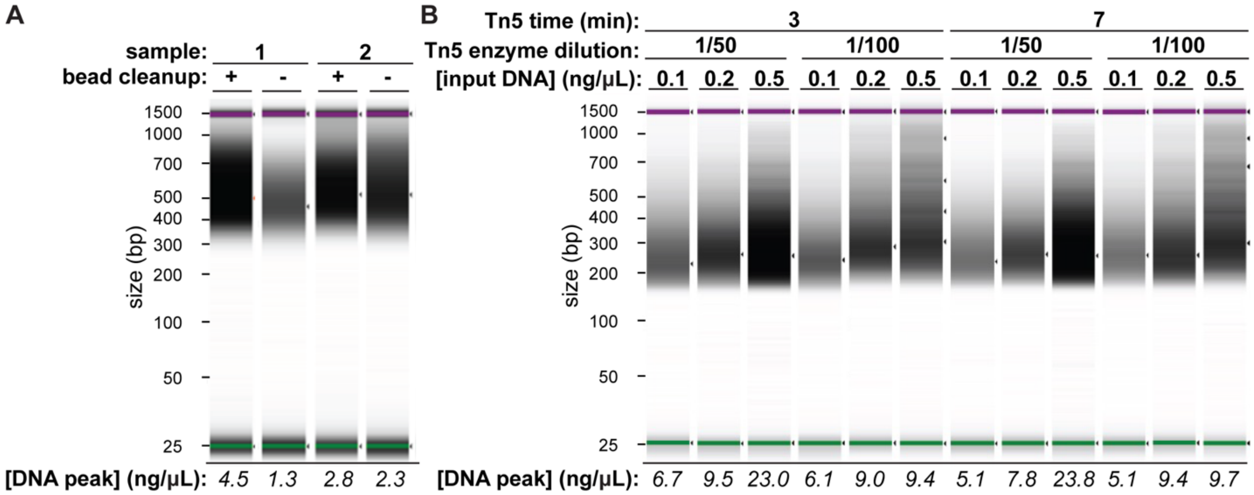

**Figure S7**. Electropherograms of tagmented and amplified mutant sublibraries.

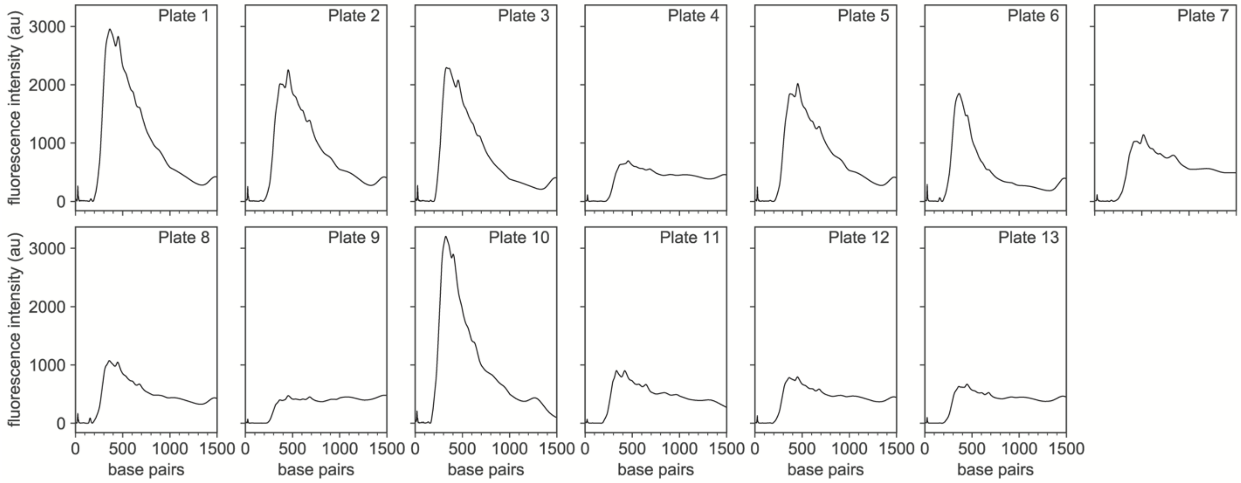

**Figure S8**. Histogram of variant:WT read ratios among single, double, and triple and greater mutants.

Read counts for variant and WT sequences represent the sum of forward and reverse reads of each genotype, averaged across each nucleotide for each codon substitution. If higher order mutants originated solely from the presence of co-occurring mutations on the same plasmid genome, the rate of WT reads for each substitution would be comparable for all types of mutants. However, a higher variant:WT read ratio for single mutants compared to double, and triple and greater is consistent with the model that many higher order mutants originate from well-to-well cross-contamination during plate handling steps prior to barcoding.

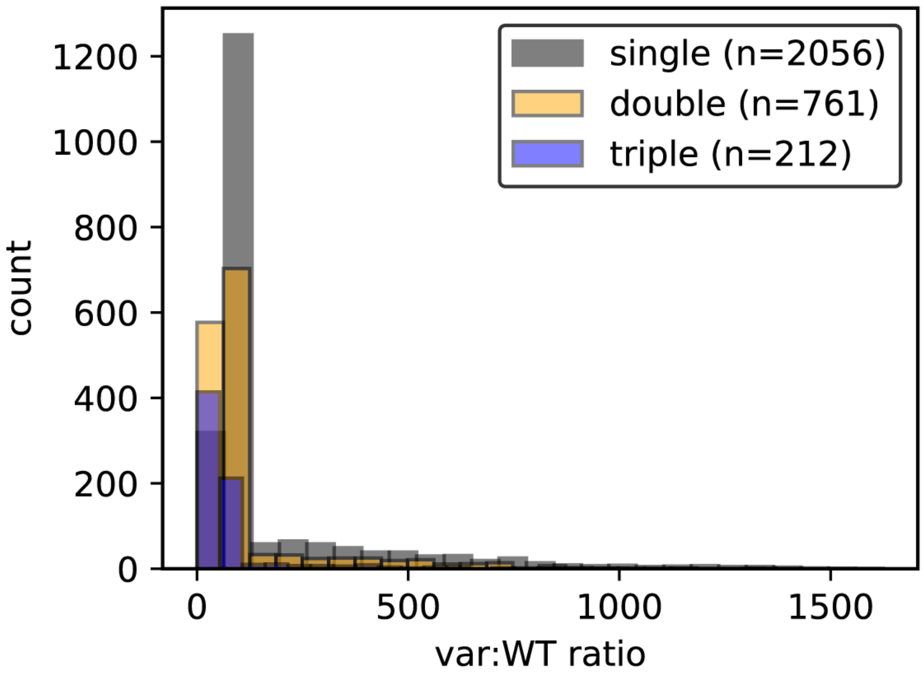

**Figure S9**. Comparison of observed and simulated single mutant frequency distributions.
We simulated predicted single mutant frequency distributions per sublibrary (assuming equal relative abundances among individual single mutant genotypes) and calculated the following three parameters: total number of barcodes (of 384 possible) meeting a read depth threshold, fraction of intended single mutants compared to all other library variants, and the number of desired unique single mutants. These parameters were used to simulate picking experiments as described in Results and Discussion, and repeated for 1000 replicates (orange bars). Simulated distributions were plotted alongside observed single mutant frequencies for each sublibrary (blue bars).

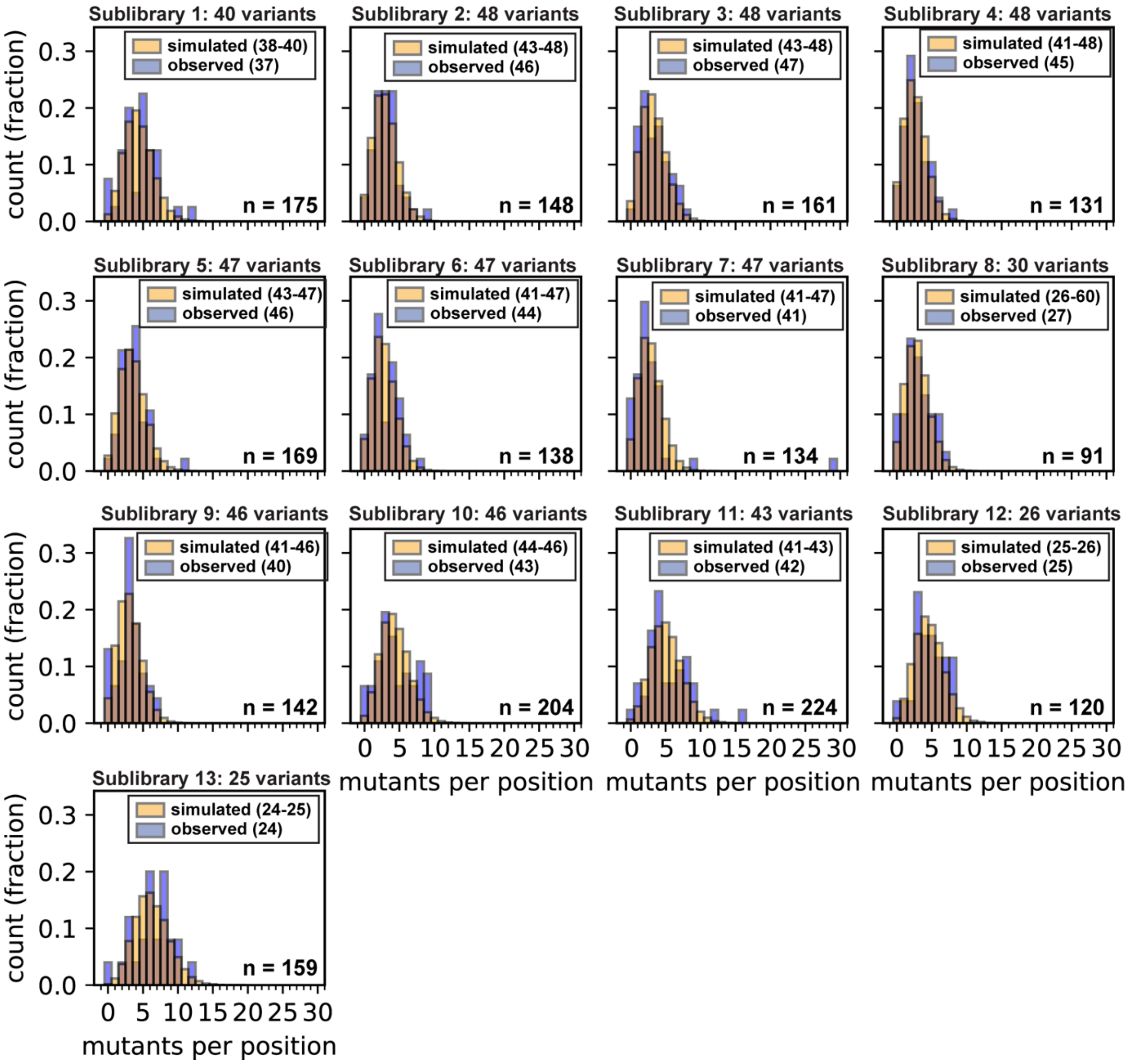

**Figure S10**. Plasmid map of PURExpress-SpAP-eGFP.

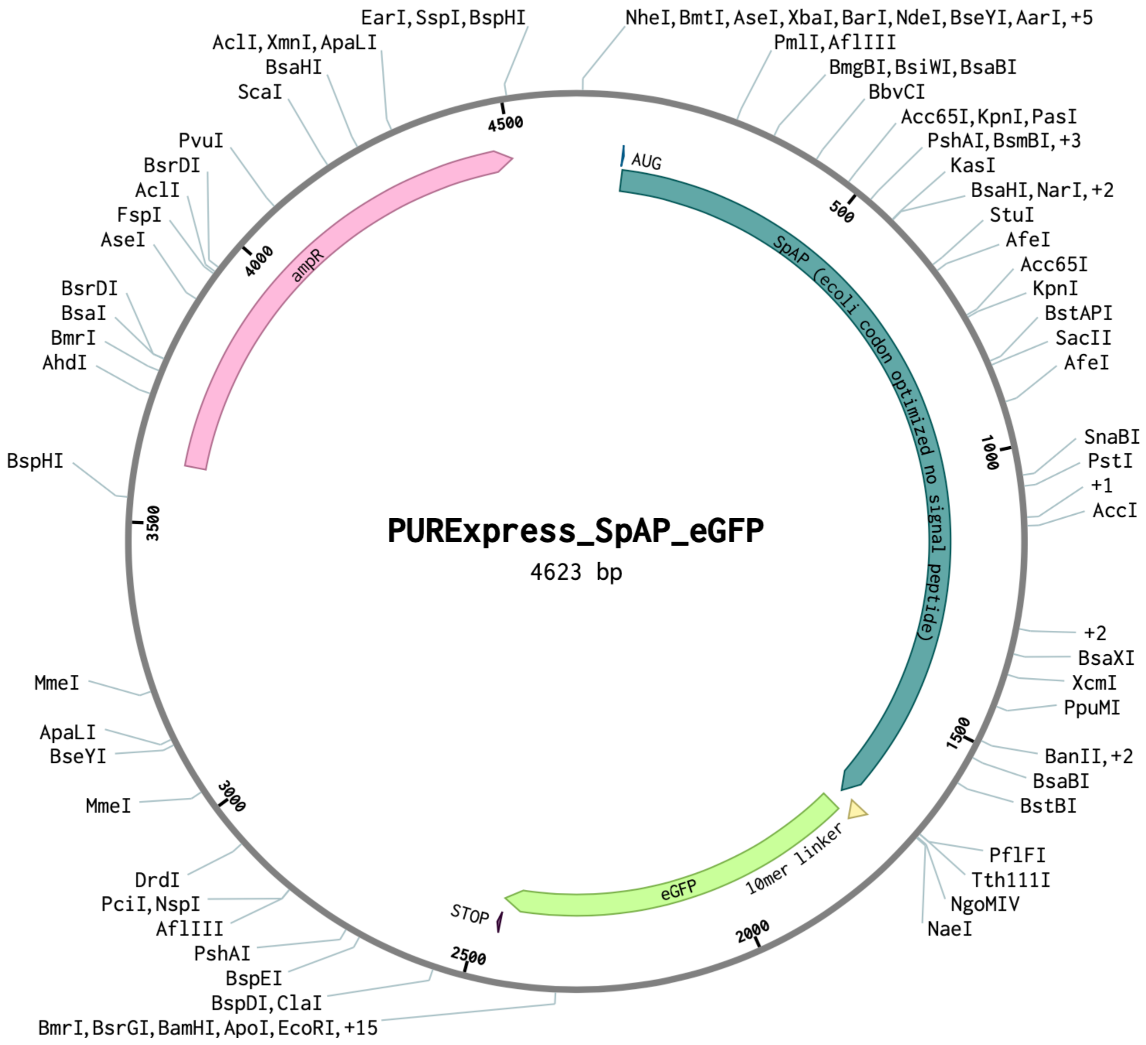

**Figure S11**. Complete DNA sequence of PURExpress-SpAP-eGFP plasmid.

GCTAGTGGTGCTAGCCCCGCGAAATTAATACGACTCACTATAGGGTCTAGAAATAATTTTGTTTAACTTTAAGAAGGAGATATAC
ATATGCAAAGCCCAGCACCTGCCGCAGCGCCTGCCCCTGCGGCACGTTCCATCGCAGCTACGCCTCCTAAACTGATCGTGGC
AATTAGCGTGGACCAGTTTAGTGCAGACTTGTTCTCGGAGTATCGTCAATATTACACCGGAGGTTTAAAGCGTCTTACATCCGAA
GGAGCTGTGTTCCCACGTGGTTATCAGAGTCATGCGGCAACAGAAACGTGTCCTGGTCACTCAACGATCCTGACAGGATCACG
TCCGTCACGTACGGGTATTATCGCTAATAACTGGTTCGACTTGGACGCAAAGCGTGAGGATAAAAATCTGTACTGTGCTGAGGA
TGAATCCCAACCCGGTAGTTCGTCTGACAAGTACGAAGCTTCGCCACTGCACTTAAAGGTACCCACCCTGGGGGGACGCATGA
AAGCCGCCAATCCTGCGACTCGTGTCGTCTCTGTTGCCGGCAAGGATCGCGCGGCCATTATGATGGGTGGCGCCACAGCGGA
TCAGGTCTGGTGGTTAGGGGGGCCTCAGGGGTATGTTTCGTATAAGGGTGTAGCGCCAACTCCCCTTGTAACACAGGTCAATC
AGGCCTTTGCACAGCGCTTAGCTCAGCCGAACCCGGGATTTGAGTTGCCTGCTCAGTGCGTCAGCAAGGACTTTCCTGTTCAA
GCGGGAAATCGCACAGTGGGTACCGGCCGCTTCGCCCGTGATGCTGGTGACTACAAAGGTTTTCGCATTTCCCCGGAGCAGG
ATGCTATGACGCTTGCATTCGCTGCCGCGGCCATTGAAAATATGCAATTAGGGAAGCAGGCCCAGACCGATATTATTAGCATTG
GACTGAGCGCTACGGATTACGTGGGACACACCTTCGGCACGGAGGGTACGGAGAGTTGCATCCAAGTGGATCGTTTAGACAC
GGAGCTTGGTGCATTCTTTGATAAACTGGATAAGGATGGGATTGACTACGTAGTAGTGCTGACTGCAGATCATGGAGGACACGA
TCTGCCCGAACGTCATCGTATGAATGCCATGCCGATGGAACAGCGCGTAGACATGGCCCTGACACCTAAAGCTCTGAATGCTA
CCATCGCTGAGAAAGCTGGCCTTCCGGGCAAAAAGGTTATTTGGTCAGATGGACCTTCTGGCGATATTTACTATGATAAGGGCC
TTACAGCCGCTCAACGTGCCCGTGTTGAAACCGAGGCGTTAAAATACTTGCGCGCGCATCCCCAAGTACAGACTGTATTCACTA
AGGCGGAAATCGCGGCTACCCCTTCTCCGTCGGGACCACCTGAGAGCTGGAGTTTGATCCAGGAAGCTCGCGCGTCATTTTAC
CCGTCGCGCTCCGGGGACCTGTTACTTTTATTGAAACCTCGTGTGATGAGCATTCCTGAGCAAGCAGTCATGGGCTCGGTTGC
AACCCATGGATCTCCATGGGATACGGATCGCCGTGTGCCTATCCTGTTTTGGCGCAAAGGTATGCAGCATTTCGAACAACCCTT
AGGAGTAGAGACTGTTGATATTTTGCCCTCCTTGGCTGCACTTATTAAGCTTCCTGTTCCTAAGGATCAGATCGACGGCCGCTG
TCTGGACTTGGTCGCCGGCAAGGATGATTCCTGTGCTGGACAGGGAGGAGGGTCTGGGGGAGGAGGCAGTGGCATGGTGAG
CAAGGGCGAGGAGCTGTTCACCGGGGTGGTGCCCATCCTGGTCGAGCTGGACGGCGACGTAAACGGCCACAAGTTCAGCGT
GTCCGGCGAGGGCGAGGGCGATGCCACCTACGGCAAGCTGACCCTGAAGTTCATCTGCACCACCGGCAAGCTGCCCGTGCC
CTGGCCCACCCTCGTGACCACCCTGACCTACGGCGTGCAGTGCTTCAGCCGCTACCCCGACCACATGAAGCAGCACGACTTC
TTCAAGTCCGCCATGCCCGAAGGCTACGTCCAGGAGCGCACCATCTTCTTCAAGGACGACGGCAACTACAAGACCCGCGCCG
AGGTGAAGTTCGAGGGCGACACCCTGGTGAACCGCATCGAGCTGAAGGGCATCGACTTCAAGGAGGACGGCAACATCCTGGG
GCACAAGCTGGAGTACAACTACAACAGCCACAACGTCTATATCATGGCCGACAAGCAGAAGAACGGCATCAAGGTGAACTTCA
AGATCCGCCACAACATCGAGGACGGCAGCGTGCAGCTCGCCGACCACTACCAGCAGAACACCCCCATCGGCGACGGCCCCG
TGCTGCTGCCCGACAACCACTACCTGAGCACCCAGTCCGCCCTGAGCAAAGACCCCAACGAGAAGCGCGATCACATGGTCCT
GCTGGAGTTCGTGACCGCCGCCGGGATCACTCTCGGCATGGACGAGCTGTACAAATAATAATGAGGATCCCGGGAATTCTCGA
GTAAGGTTAACCTGCAGGAGGCCTTTAATTAAGGTGGTGCGGCCGCGCTAGCGGTCCCGGGGGATCGATCCGGCTGCTAACA
AAGCCCGAAAGGAAGCTGAGTTGGCTGCTGCCACCGCTGAGCAATAACTAGCATAACCCCTTGGGGCCTCTAAACGGGTCTTG
AGGGGTTTTTTGCTGAAAGGAGGAACTATATCCGGAAGCTTGGCACTGGCCGACCGGGGTCGAGCACTGACTCGCTGCGCTC
GGTCGTTCGGCTGCGGCGAGCGGTATCAGCTCACTCAAAGGCGGTAATACGGTTATCCACAGAATCAGGGGATAACGCAGGA
AAGAACATGTGAGCAAAAGGCCAGCAAAAGGCCAGGAACCGTAAAAAGGCCGCGTTGCTGGCGTTTTTCCATAGGCTCCGCCC
CCCTGACGAGCATCACAAAAATCGACGCTCAAGTCAGAGGTGGCGAAACCCGACAGGACTATAAAGATACCAGGCGTTTCCCC
CTGGAAGCTCCCTCGTGCGCTCTCCTGTTCCGACCCTGCCGCTTACCGGATACCTGTCCGCCTTTCTCCCTTCGGGAAGCGTG
GCGCTTTCTCATAGCTCACGCTGTAGGTATCTCAGTTCGGTGTAGGTCGTTCGCTCCAAGCTGGGCTGTGTGCACGAACCCCC
CGTTCAGCCCGACCGCTGCGCCTTATCCGGTAACTATCGTCTTGAGTCCAACCCGCTAAGACACGACTTATCGCCACTGGCAG
CAGCCACTGGTAACAGGATTAGCAGAGCGAGGTATGTAGGCGGTGCTACAGAGTTCTTGAAGTGGTGGCCTAACTACGGCTAC
ACTAGAAGAACAGTATTTGGTATCTGCGCTCTGCTGAAGCCAGTTACCTTCGGAAAAAGAGTTGGTAGCTCTTGATCCGGCAAA
CAAACCACCGCTGGTAGCGGTGGTTTTTTTGTTTGCAAGCAGCAGATTACGCGCAGAAAAAAAGGATCTCAAGAAGATCCTTTG
ATCTTTTCTACGGGGTCTGACGCTCAGTGGAACGAAAACTCACAGATCCGGGATTTTGGTCATGAGATTATCAAAAAGGATCTT
CACCTAGATCCTTTTAAATTAAAAATGAAGTTTTAAATCAATCTAAAGTATATATGAGTAAACTTGGTCTGACAGTTACCAATGCTT
AATCAGTGAGGCACCTATCTCAGCGATCTGTCTATTTCGTTCATCCATAGTTGCCTGACTCCCCGTCGTGTAGATAACTACGATA
CGGGAGGGCTTACCATCTGGCCCCAGTGCTGCAATGATACCGCGAGACCCACGCTCACCGGCTCCAGATTTATCAGCAATAAA
CCAGCCAGCCGGAAGGGCCGAGCGCAGAAGTGGTCCTGCAACTTTATCCGCCTCCATCCAGTCTATTAATTGTTGCCGGGAAG
CTAGAGTAAGTAGTTCGCCAGTTAATAGTTTGCGCAACGTTGTTGCCATTGCTACAGGCATCGTGGTGTCACGCTCGTCGTTTG
GTATGGCTTCATTCAGCTCCGGTTCCCAACGATCAAGGCGAGTTACATGATCCCCCATGTTGTGCAAAAAAGCGGTTAGCTCCT
TCGGTCCTCCGATCGTTGTCAGAAGTAAGTTGGCCGCAGTGTTATCACTCATGGTTATGGCAGCACTGCATAATTCTCTTACTGT
CATGCCATCCGTAAGATGCTTTTCTGTGACTGGTGAGTACTCAACCAAGTCATTCTGAGAATAGTGTATGCGGCGACCGAGTTG
CTCTTGCCCGGCGTCAATACGGGATAATACCGCGCCACATAGCAGAACTTTAAAAGTGCTCATCATTGGAAAACGTTCTTCGGG
GCGAAAACTCTCAAGGATCTTACCGCTGTTGAGATCCAGTTCGATGTAACCCACTCGTGCACCCAACTGATCTTCAGCATCTTTT
ACTTTCACCAGCGTTTCTGGGTGAGCAAAAACAGGAAGGCAAAATGCCGCAAAAAAGGGAATAAGGGCGACACGGAAATGTTG
AATACTCATACTCTTCCTTTTTCAATATTATTGAAGCATTTATCAGGGTTATTGTCTCATGAGCGGATACATATTTGAATGTATTTA
GAAAAATAAACAAATAGGGGTTCCGCGCACATTTCCCCGAAAAGT

**Figure S12**. Protein sequence of SpAP-(10mer linker)-eGFP.
MQSPAPAAAPAPAARSIAATPPKLIVAISVDQFSADLFSEYRQYYTGGLKRLTSEGAVFPRGYQ
SHAATETCPGHSTILTGSRPSRTGIIANNWFDLDAKREDKNLYCAEDESQPGSSSDKYEASPLH
LKVPTLGGRMKAANPATRVVSVAGKDRAAIMMGGATADQVWWLGGPQGYVSYKGVAPTPLV
TQVNQAFAQRLAQPNPGFELPAQCVSKDFPVQAGNRTVGTGRFARDAGDYKGFRISPEQDA
MTLAFAAAAIENMQLGKQAQTDIISIGLSATDYVGHTFGTEGTESCIQVDRLDTELGAFFDKLDK
DGIDYVVVLTADHGGHDLPERHRMNAMPMEQRVDMALTPKALNATIAEKAGLPGKKVIWSDG
PSGDIYYDKGLTAAQRARVETEALKYLRAHPQVQTVFTKAEIAATPSPSGPPESWSLIQEARAS
FYPSRSGDLLLLLKPRVMSIPEQAVMGSVATHGSPWDTDRRVPILFWRKGMQHFEQPLGVET
VDILPSLAALIKLPVPKDQIDGRCLDLVAGKDDSCAGQGGGSGGGGSGMVSKGEELFTGVVPIL
VELDGDVNGHKFSVSGEGEGDATYGKLTLKFICTTGKLPVPWPTLVTTLTYGVQCFSRYPDHM
KQHDFFKSAMPEGYVQERTIFFKDDGNYKTRAEVKFEGDTLVNRIELKGIDFKEDGNILGHKLE
YNYNSHNVYIMADKQKNGIKVNFKIRHNIEDGSVQLADHYQQNTPIGDGPVLLPDNHYLSTQSA
LSKDPNEKRDHMVLLEFVTAAGITLGMDELYK

**Table S1**. Oligo array and window design details for SpAP scanning mutant library.

| sublibrary | length (nt) | primers (forward, reverse) | SpAP positions mutated | substitution |
|---|---|---|---|---|
| 1 | 200 | CGACTCACTATAGGGTCTAGAAATA, CCTCCGGTGTAATATTGACG | 2–41 | non-val → val<br>val → ala |
| 2 | 200 | GTTTAGTGCAGACTTGTTCTCGGAGT, CGTCCAAGTCGAACCAGTTATTAGCG | 42–89 | non-val → val<br>val → ala |
| 3 | 197 | CTGACAGGATCACGTCCGTCAC, GCAGGATTGGCGGCTTTCAT | 90–137 | non-val → val<br>val → ala |
| 4 | 199 | CTTAAAGGTACCCACCCTGGGG, CTGATTGACCTGTGTTACAAGGGGAG | 138–185 | non-val → val<br>val → ala |
| 5 | 197 | GGGGTATGTTTCGTATAAGGGTGTAGC, CTTTGTAGTCACCAGCATCACGGGC | 186–232 | non-val → val<br>val → ala |
| 6 | 196 | CGGGAAATCGCACAGTGGGTAC, GTGTGTCCCACGTAATCCGTAGC | 233–279 | non-val → val<br>val → ala |
| 7 | 197 | GCCCAGACCGATATTATTAGCATTGGAC, GGCAGATCGTGTCCTCCATGATC | 280–326 | non-val → val<br>val → ala |
| 8 | 193 | CGTTTAGACACGGAGCTTGGTG, CTTTCTCAGCGATGGTAGCATTCAG | 327–356 | non-ala → ala<br>ala → ala<br>(synonymous) |
| 9 | 193 | GACATGGCCCTGACACCTAAAGC, CAGTCTGTACTTGGGGATGCGC | 357–402 | non-ala → ala<br>ala → ala<br>(synonymous) |
| 10 | 199 | CAACGTGCCCGTGTTGAAACC, CACGAGGTTTCAATAAAAGTAACAGGTC | 403–448 | non-ala → ala<br>ala → ala<br>(synonymous) |
| 11 | 181 | CGTCATTTTACCCGTCGCGCTC, CCTAAGGGTTGTTCGAAATGCTGC | 449–491 | non-ala → ala<br>ala → ala<br>(synonymous) |
| 12 | 134 | GCCGTGTGCCTATCCTGTTTTG, GCCGTCGATCTGATCCTTAGGAAC | 492–517 | non-ala → ala<br>ala → ala<br>(synonymous) |
| 13[a] | 139 | CCTTGGCTGCACTTATTAAGCTTCC, TTGCTCACCATGCCACTGCCTC | 518–542 | non-ala → ala<br>ala → ala<br>(synonymous) |

[a]This sublibrary also encodes a mutation for position 542, which is the first residue within a 10 amino acid linker between the SpAP and eGFP ORFs.

**Table S2**. Concentration of purified sublibrary mutagenic primer pools

| sublibrary | primer pool concentration (nM)[a] |
|---|---|
| 1 | 17.5 |
| 2 | 28.8 |
| 3 | 23.8 |
| 4 | 31.6 |
| 5 | 51.2 |
| 6 | 37.2 |
| 7 | 20.4 |
| 8 | 14.7 |
| 9 | 27.6 |
| 10 | 40.0 |
| 11 | 61.2 |
| 12 | 19.1 |
| 13 | 34.0 |

[a]Purified dsDNA samples were quantified by UV absorbance.

**Table S3**. Expected mutant yields from simulations of mutant sampling.
Sequenced clones are reported for achieving unique mutant yields equivalent to 90% of total mutants.

| total mutants | single mutant frequency | sequenced clones (median)[a] | sequenced clones (lower)[b] | sequenced clones (upper)[c] |
|---|---|---|---|---|
| 50 | 0.10 | >500 | >500 | >500 |
| 50 | 0.25 | 445 | 324 | >500 |
| 50 | 0.50 | 218 | 162 | 473 |
| 50 | 0.75 | 144 | 105 | 450 |
| 50 | 1.00 | 106 | 85 | 330 |
| 500 | 0.10 | >5000 | >5000 | >5000 |
| 500 | 0.25 | 4726 | 4290 | >5000 |
| 500 | 0.50 | 2322 | 2081 | 4793 |
| 500 | 0.75 | 1538 | 1407 | 4702 |
| 500 | 1.00 | 1147 | 1056 | 4309 |
| 5000 | 0.10 | >50000 | >50000 | >50000 |
| 5000 | 0.25 | 47703 | 46405 | 49957 |
| 5000 | 0.50 | 23431 | 22589 | 49832 |
| 5000 | 0.75 | 15503 | 15080 | 49446 |
| 5000 | 1.00 | 11586 | 11296 | 46054 |

[a]The minimum number of sequenced clones required to obtain 90% yield of unique single mutants, as determined by the median unique mutant yield of 100 simulated picking experiments.

[b]The lower bound was calculated as the minimum number of sequenced clones required to obtain a 90% yield of unique single mutants within the 95% confidence interval of unique mutant yields expected for this volume, from 100 simulated picking experiments.

[c]The upper bound was calculated as the maximum number of sequenced clones required to obtain a 90% yield of unique single mutants within the 95% confidence interval of unique mutant yields expected for this volume, from 100 simulated picking experiments.

**Table S4**. Variant composition of small-scale QuikChange-HT reactions.

| **variant type**[a] | **count (n=96)** | **fraction** |
|---|---|---|
| WT | 11 | 0.11 |
| single | 60 | 0.63 |
| double | 4 | 0.04 |
| triple+ | 1 | 0.01 |
| indels, errors[b] | 20 | 0.21 |

[a]Clones were sequenced with one forward primer spanning the mutational region
[b]Includes indels, with or without the presence of intended codon substitution(s), and includes errors likely attributable to sanger sequencing

**Table S5**. Sublibrary transformation and colony picking results.

| sublibrary | primer pool concentration (nM) | colonies[a] | repeat QuikChange[b] |
|---|---|---|---|
| 1 | 17.5 | 76 | Y |
| 2 | 28.8 | 136 | N |
| 3 | 23.8 | 130 | N |
| 4 | 31.6 | 200 | N |
| 5 | 51.2 | 440 | N |
| 6 | 37.2 | 25 | Y |
| 7 | 20.4 | 26 | Y |
| 8 | 14.7 | 130 | N |
| 9 | 27.6 | 220 | N |
| 10 | 40.0 | 80 | N |
| 11 | 61.2 | 150 | N |
| 12 | 19.1 | 120 | N |
| 13 | 34.0 | 220 | N |

[a]Number of colonies obtained after QuikChange mutagenesis using normalized primer concentrations of 15 nM. Plating details: 1 µL of reaction volume used to transform 20 µL NEB-5α cells, followed by addition of 200 µL SOC, of which 100 µL was plated on a 150 mm LB agar plate. Sublibraries not meeting colony yield requirements (400–500 colonies) were plated and/or transformed again at higher volume.

[b]QuikChange mutagenesis was repeated for these sublibraries using the maximum possible concentrations of stock primer pools allowed by reaction volumes.

**Table S6**. Amplicon DNA and library concentration statistics.

| sublibrary | plate | sample wells total | sample wells assayed[a] | median[b] | mean[b] | standard deviation[b] | tagmented library concentration[b, c] |
|---|---|---|---|---|---|---|---|
| 1 | 1 | 384 | 184 | 25 | 34 | 24 | 17 |
| 2 | 2 | 384 | 184 | 61 | 58 | 22 | 12 |
| 3 | 3 | 384 | 184 | 37 | 39 | 17 | 14 |
| 4 | 4 | 384 | 184 | 28 | 32 | 22 | 3 |
| 5 | 5 | 384 | 184 | 39 | 40 | 26 | 12 |
| 6 | 6 | 384 | 184 | 11 | 11 | 7 | 9 |
| 7 | 7 | 384 | 184 | 29 | 32 | 21 | 4 |
| 8 | 8 | 384 | 368 | 1 | 20 | 27 | 5 |
| 9 | 9 | 384 | 184 | 47 | 47 | 23 | 3 |
| 10 | 10 | 384 | 184 | 57 | 54 | 29 | 17 |
| 11 | 11 | 384 | 184 | 31 | 34 | 21 | 3 |
| 12 | 12 | 384 | 184 | 16 | 18 | 11 | 4 |
| 13 | 13 | 384 | 184 | 17 | 21 | 14 | 3 |

[a]Number of amplicon wells measured by fluorescence assay for DNA concentration

[b]In units of ng/μL

[c]Following tagmentation, barcoding/amplification PCR, and pooling of all 384 sample wells per plate, concentration represents total upon integration of all fragmentation peaks (see Figure 5)

**Table S7**. Unique single mutant yields for the SpAP scanning library.
Total and fractional yields for the entire library (bold text) and within each mutational sublibrary are shown at read threshold values of 0, 1, 10, 100, and 1000. The read threshold value represents the minimum number of variant reads for each single mutant, and, the minimum ratio of var:WT reads (sum of forward and reverse reads in each case).

| sublibrary | residues | total positions | yield at read thresholds: 0–1000 (fraction of total) | | | |
|---|---|---|---|---|---|---|
| | | | n/a | 10 | 100 | 1000 |
| **all** | **2–542** | **541** | **507 (0.94)** | **498 (0.92)** | **484 (0.89)** | **60 (0.11)** |
| 1 | 2–41 | 40 | 37 (0.93) | 36 (0.9) | 35 (0.88) | 0 (0) |
| 2 | 42–89 | 48 | 46 (0.96) | 45 (0.94) | 45 (0.94) | 12 (0.25) |
| 3 | 90–137 | 48 | 47 (0.98) | 46 (0.96) | 45 (0.94) | 7 (0.15) |
| 4 | 138–185 | 48 | 45 (0.94) | 41 (0.85) | 36 (0.75) | 2 (0.04) |
| 5 | 186–232 | 47 | 46 (0.98) | 46 (0.98) | 45 (0.96) | 6 (0.13) |
| 6 | 233–279 | 47 | 44 (0.94) | 44 (0.94) | 42 (0.89) | 1 (0.02) |
| 7 | 280–326 | 47 | 41 (0.87) | 40 (0.85) | 40 (0.85) | 3 (0.06) |
| 8 | 327–356 | 30 | 27 (0.9) | 26 (0.87) | 26 (0.87) | 4 (0.13) |
| 9 | 357–402 | 46 | 40 (0.87) | 40 (0.87) | 38 (0.83) | 1 (0.02) |
| 10 | 403–448 | 46 | 43 (0.93) | 43 (0.93) | 42 (0.91) | 8 (0.17) |
| 11 | 449–491 | 43 | 42 (0.98) | 42 (0.98) | 42 (0.98) | 16 (0.37) |
| 12 | 492–517 | 26 | 25 (0.96) | 25 (0.96) | 24 (0.92) | 0 (0) |
| 13 | 518–542 | 25 | 24 (0.96) | 24 (0.96) | 24 (0.96) | 0 (0) |

**Table S8**. Comparison of uPIC–M performance with simulated picking experiments, per sublibrary.

| sublibrary | total barcodes | single mutant frequency | total positions | observed positions[a] | expected positions, median (95% CI)[b] |
|---|---|---|---|---|---|
| 1 | 318 | 0.55 | 40 | **37** | 40 (38–40) |
| 2 | 274 | 0.54 | 48 | 46 | 46 (43–48) |
| 3 | 298 | 0.54 | 48 | 47 | 46 (43–48) |
| 4 | 247 | 0.53 | 48 | 45 | 45 (41–48) |
| 5 | 315 | 0.54 | 47 | 46 | 46 (43–47) |
| 6 | 264 | 0.52 | 47 | 44 | 45 (41–47) |
| 7 | 241 | 0.56 | 47 | 41 | 45 (41–47) |
| 8 | 146 | 0.62 | 30 | 27 | 29 (26–30) |
| 9 | 260 | 0.55 | 46 | **40** | 44 (41–46) |
| 10 | 352 | 0.58 | 46 | **43** | 46 (44–46) |
| 11 | 329 | 0.68 | 43 | 42 | 43 (41–43) |
| 12 | 216 | 0.56 | 26 | 25 | 26 (25–26) |
| 13 | 270 | 0.59 | 25 | 24 | 25 (24–25) |

[a]Number of unique mutants obtained for sublibrary. Sublibraries not meeting expected yields (based on the 95% confidence interval) are in bold red text.

[b]Predicted number of unique mutants given the observed single mutant frequency and number of clones sampled for plate, reported as the median of 1000 simulated sampling events.

**Table S9**. Oligo array price summary.

| array size (unique oligos) | array cost (USD) | cost per oligo (USD)[a] |
|---|---|---|
| $7.5 \times 10^3$ | 2,857 | 0.38 |
| $1.5 \times 10^4$ | 5,714 | 0.38 |
| $6.0 \times 10^4$ | 8,435 | 0.14 |
| $1.0 \times 10^5$ | 10,856 | 0.11 |
| $2.44 \times 10^5$ | 23,883 | 0.10 |

[a]Oligo array price summary (academic pricing, obtained 01/2021 from Agilent Technologies, personal communication). Prices are provided for arrays containing oligos of length 191–210 nt.
